# A home and rescue gene drive efficiently spreads and persists in populations

**DOI:** 10.1101/2020.08.21.261610

**Authors:** Nikolay P. Kandul, Junru Liu, Jared B. Bennett, John M. Marshall, Omar S. Akbari

## Abstract

Homing based gene drives, engineered using CRISPR/Cas9, have been proposed to spread desirable genes into target populations. However, spread of such drives can be hindered by the accumulation of resistance alleles. To overcome this significant obstacle, we engineer an inherently confinable population modification <u>Home</u>-and-<u>R</u>escue (HomeR) drive in *Drosophila melanogaster* that, by creative design, limits the accumulation of such alleles. We demonstrate that HomeR can achieve nearly ∼100% transmission enabling it to spread and persist at genotypic fixation in several multi-generational population cage experiments, underscoring its long term stability and drive potential. Finally, we conduct mathematical modeling determining HomeR can outperform contemporary gene drive architectures for population modification over wide ranges of fitness and transmission rates. Given its straightforward design, HomeR could be universally adapted to a wide range of species.

## Introduction

Effective insect control strategies are necessary for preventing human diseases, such as malaria and dengue virus, and protecting crops from pests. These challenges have fostered the development of innovative population control technologies such as Cas9/guideRNA (Cas9/gRNA) homing-based gene drives (HGDs) (Champer et al., 2016; Esvelt et al., 2014) which have been tested in the laboratory for either population modification (Adolfi et al., 2020; Carballar-Lejarazú et al., 2020; Gantz et al., 2015; Li et al., 2020; Pham et al., 2019) to spread desirable traits that can impair the mosquitoes ability to transmit pathogens (e.g. (Buchman et al., 2020, 2019; Hoermann et al., 2020; Isaacs et al., 2012; Marshall et al., 2019)), or population suppression (Hammond et al., 2016; Kyrou et al., 2018; Simoni et al., 2020) to reduce and eliminate wild disease transmitting populations of mosquitoes. Despite significant progress, HGDs are still an emerging technology that can suffer from the formation of resistance alleles, hindering their efficacy (Adolfi et al., 2020; Carballar-Lejarazú et al., 2020; Gantz et al., 2015; Hammond et al., 2016; Kandul et al., 2019a; Kyrou et al., 2018; Li et al., 2020; Pham et al., 2019; Simoni et al., 2020).

In CRISPR/Cas9, the Cas9 endonuclease cuts a programmed DNA sequence complementary to a user defined short guide RNA molecule (gRNA). To engineer a HGD, CRISPR components are integrated at the cut site in the genome so that when they cut the recipient wildtype (*wt*) allele it is repaired via homology-directed repair (HDR) in heterozygotes, using the donor allele (i.e. allele harboring the HGD) as a reference for DNA repair. This enables the HGD to home, or copy, itself into the recipient allele (Champer et al., 2016; Esvelt et al., 2014) (referred to as homing from hereon). This general design for HGD was quickly adopted, and many HGDs were developed in several insect species (Gantz et al., 2015; Hammond et al., 2016; Kandul et al., 2019a; Kyrou et al., 2018; Li et al., 2020; Simoni et al., 2020). However, it soon became apparent that HGDs unintentionally promote the formation of resistance alleles through mutagenic repair. When these alleles are positively selected they can hinder HGD spread in laboratory cage populations (Champer et al., 2017; Hammond et al., 2017; Kandul et al., 2019a; KaramiNejadRanjbar et al., 2018; Oberhofer et al., 2018), with one exception that targeted an ultra-conserved sex determination gene for population suppression (Kyrou et al., 2018; Simoni et al., 2020). This resistance arises from, in addition to HDR, Cas9/gRNA-directed DNA cuts are also repaired by non-homologous end joining (NHEJ), an alternative DNA repair pathway that occasionally introduces insertions or deletions (*indels*) at the target site. Many of these indels produce loss-of-function (LOF) alleles, which can be selected against. However, functional in-frame NHEJ-induced *indel* alleles can propagate, that are unrecognized by the same Cas9/gRNA complex, and become drive resistance alleles. When resistance alleles are induced in germ cells, they are heritable and can hinder spread of HGDs (Champer et al., 2017; Hammond et al., 2017; Kandul et al., 2019a; KaramiNejadRanjbar et al., 2018; Oberhofer et al., 2018). Both induced and naturally existing resistance alleles can pose serious challenges to engineering a stable HGD capable of spreading and persisting in a population.

To overcome the accumulation of drive resistance alleles, CRISPR based toxin-antidote (TA) based systems, in which embryos are essentially “poisoned” and only those embryos harboring the TA genetic cassette are rescued, were described (Fig. S8 in (Kandul et al., 2019b)) and engineered (Champer et al., 2020a; Oberhofer et al., 2020a, 2020b, 2019). Generally these designs utilize a toxin consisting of a non-homing GD harboring multiple gRNAs targeting a vital gene, and an “addictive” antidote that is a re-coded, cleavage-immune version of the targeted gene. These TA based drives are Mendelianly transmitted and spread by killing progeny that fail to inherit the drive (e.g. 50% perish from heterozygous mother). Alternative HDR-based TA designs were also described (Champer et al., 2016; Esvelt et al., 2014), modeled (Noble et al., 2017), and recently tested in mosquitoes (Adolfi et al., 2020) targeting recessive non-essential genes for viability, and in *Drosophila melanogaster* targeting a rare haploinsufficient (i.e. the non-functional allele is dominant as a single functional copy of the target gene is <u>not sufficient</u> for normal function) gene (Champer et al., 2020b), each demonstrating drive capacity.

Building upon prior work, here we describe the development of a <u>Home</u>-and-<u>R</u>escue (HomeR) split-drive (i.e. Cas9 separated from the drive) targeting an essential, haplosufficient (i.e. the non-functional allele is recessive as a single functional copy of the target gene <u>is sufficient</u> for normal function) gene in *Drosophila melanogaster*. We demonstrate that the accumulation of NHEJ-induced resistance alleles can be reduced by strategically (i) designing the HGD to target the conserved 3’ coding sequence of a haplosufficient gene required for insect viability, and (ii) encoding a dominant rescue of the endogenous target gene into HomeR, and (iii) using an exogenous 3’ UTR to prevent expected deleterious recombination events between the drive and the endogenous target gene. We demonstrate that efficient cleavage of the target sequence by HomeR and rescue of the gene’s function are requisites to achieve nearly ∼100% transmission, which is accomplished by homing in ∼90% of *wt* alleles and destroying the remaining ∼10% from trans-heterozygous females. Further, we perform multi-generational population cage experiments demonstrating long term stability and efficient Cas9 dependent drive. Finally, we conduct comprehensive mathematical modeling to demonstrate that HomeR can outperform contemporary gene drive systems for population modification over wide ranges of fitness and transmission rates and, given the simplistic design, this system could be adapted to other species.

## Results

### Design and testing of gRNAs targeting an essential gene

To develop a HomeR-based drive, we first identified an essential haplosufficient gene to target. We chose *Pol-γ35*, required for the replication and repair of mitochondrial DNA (mtDNA) (Carrodeguas, 2000; Carrodeguas et al., 2001) and whose LOF results in lethality (Iyengar et al., 2002). Given that separate gRNAs can result in varying degrees of cleavage efficiencies (Kandul et al., 2019b), we tested two gRNAs targeting a conserved C-terminal domain of *Pol-γ35* (*gRNA#1*^*Pol-γ35*^ and *gRNA#2*^*Pol-γ35*^). Both *gRNA#1*^*Pol-γ35*^ and *gRNA#2*^*Pol-γ35*^ were inserted site specifically in the *Drosophila* genome and were expressed using the *U6*.*3* promoter (Port et al., 2014). To genetically assess the efficiency of *Cas9/gRNA*-mediated cleavage induced by each gRNA, we separately crossed these established gRNA lines to two different Cas9 expressing lines: (i) a previously characterized ubiquitously expressing Cas9 line (Port et al., 2014) in the DNA Ligase 4 null genetic background (*Act5C-Cas9; Lig4–/–*) (Xu Zhang et al., 2014) (**Fig. 1A**), and also to (ii) a germline-enriched Cas9 driven by the *nanos* promoter (*nos-Cas9*) (Kandul et al., 2019a, 2019b) (**Fig. 1B**). As mutations in *Lig4* gene can result in decreased activity of DNA repair by the NHEJ pathway (McVey et al., 2004) we reasoned that by testing the gRNA efficiency in a *Lig4–/–* background, we may increase the penetrance of lethality phenotypes since we are targeting an essential gene that cannot be repaired efficiently by NHEJ which simplifies scoring of the gRNAs for efficacy. Given that the *Lig4* gene is located on the X chromosome, maternal *Lig4–* alleles will be inherited by all male progeny, making them hemizygous *Lig4–* mutants, while females will be heterozygous *Lig4–/+*.

**Figure 1.**
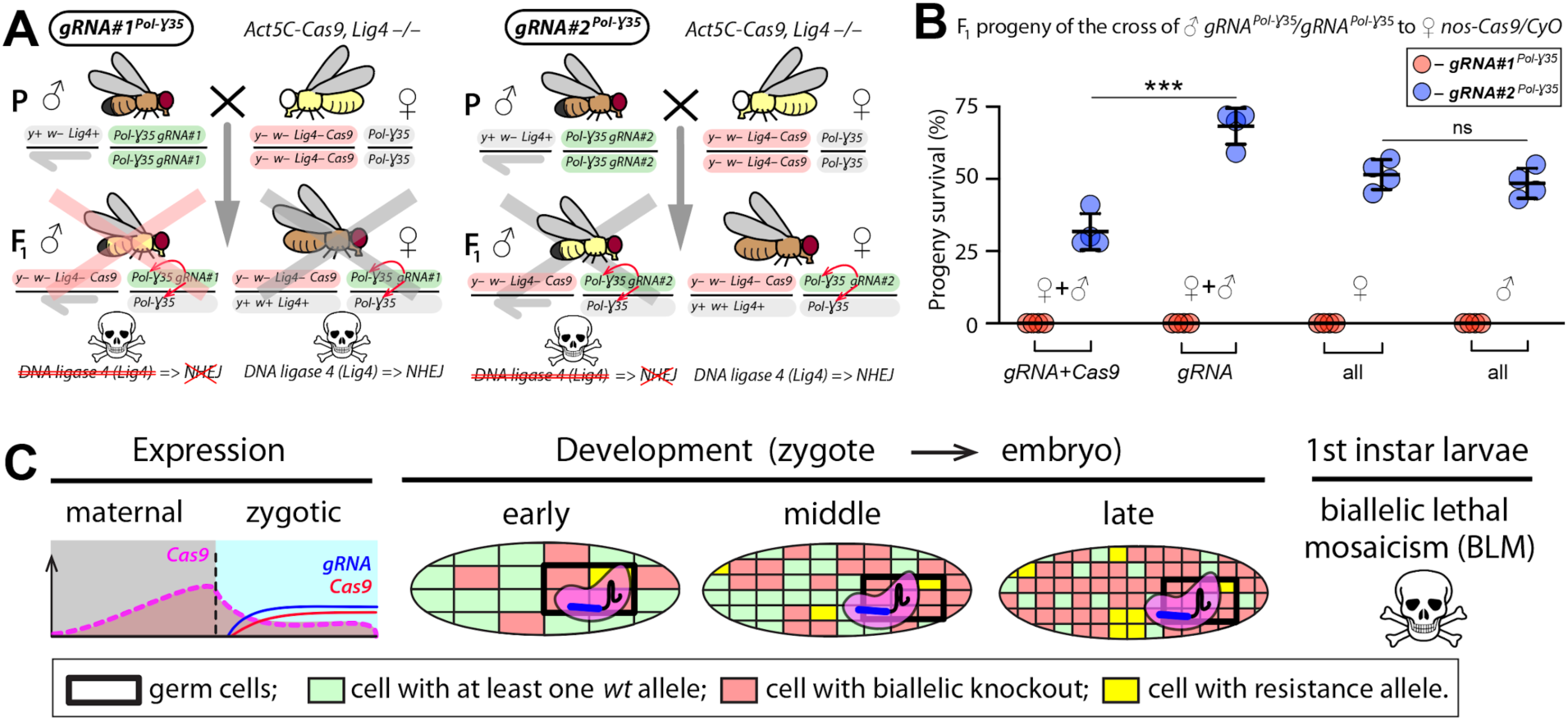
Assessing gRNAs targeting the *Drosophila Pol-γ35* gene. (**A**) Genetic crosses to assess the cleavage efficiency of two gRNAs targeting the *DNA polymerase gamma 35 kb* gene (*Pol-γ35*) in the *DNA ligase 4* null genetic background *(Lig4–/–*), in which the Non-Homologous End Joining (NHEJ) pathway is reduced. Homozygous *gRNA#1*^*Pol-γ35*^ or *gRNA#2*^*Pol-γ35*^ males were crossed to homozygous *Act5C-Cas9, Lig4 –/–* females, resulting in death of all male progeny for each gRNA. Notably, female progeny harboring *gRNA#1*^*Pol-γ35*^ and *Act5C-Cas9* in the *Lig4+/–* background also perished. (**B**) *gRNA#1*^*Pol-γ35*^, but not *gRNA#2*^*Pol-γ35*^, induced embryonic lethality of all F_1_ progeny in conjunction with *nos-Cas9* or maternal carryover of the Cas9 protein via biallelic lethal mosaicism (BLM) (Kandul et al., 2019b)(**C**). The frequency of F_1_ progeny survival presented by the genotype and/or sex. The plot shows the mean ± standard deviation (SD) over four biological replicates. Statistical significance was estimated using a two-sided Student’s *t* test with equal variance. (*p* ≥ 0.05^ns^ and *p* < 0.001***).

We observed that the genetic cross between either *U6*.*3-gRNA#1*^*Pol-γ35*^ or *U6*.*3-gRNA#2*^*Pol-γ35*^ homozygous males to *Act5C-Cas9, Lig4 –/–* homozygous females was lethal for all male progeny, presumably a result of lack of efficient repair due to the hemizygous *Lig4–* mutant background in males (**Fig. 1A**,**B**). Interestingly, trans-heterozygous *Act5C-Cas9, Lig4–/+*; *U6*.*3-gRNA#1*^*Pol-γ35*^*/+* female progeny did not survive either (**Fig. 1A**,**B, Data S1**), though they had one functional copy of the *Lig4* gene and were NHEJ proficient, suggesting that *U6*.*3-gRNA#1*^*Pol-γ35*^ is likely more potent. *U6*.*3-gRNA#1*^*Pol-γ35*^ also induced lethality in both trans-heterozygous females and males harboring *Act5C-Cas9* in the *Lig4 +/+* genetic background (**Data S2**). Furthermore, we also found that the Cas9 protein deposited by *nos-Cas9/+* females without inheritance of the *nos-Cas9* transgene, aka maternal carryover (Kandul et al., 2019b; Lin and Potter, 2016), was sufficient to ensure lethality of the F_1_ progeny harboring *U6*.*3-gRNA#1*^*Pol-γ35*^ (**Fig. 1C**), while *U6*.*3-gRNA#2*^*Pol-γ35*^ induced lethality only in a fraction of the F_1_ *U6*.*3-gRNA#2*^*Pol-γ35*^/*nos-Cas9* trans-heterozygous flies, independent of sex (**Fig. 1C, Data S2**). Pooled embryo Sanger sequencing of trans-heterozygotes revealed expected mutations at the *Pol-γ35* gRNA target sites. As we previously described, the mechanism ensuring lethality results from a dominant process we termed biallelic lethal mosaicism (BLM) (Kandul et al., 2019b), in which maternal carryover /zygotic expression results in mosaic target gene cleavage throughout development leading to wide scale loss of target gene function (**Fig. 1C**). Taken together, these results indicate that both tested gRNAs induced cleavage of the *Pol-γ35* target sequences, though *U6*.*3-gRNA#1*^*Pol-γ35*^ induced greater cleavage than *U6*.*3-gRNA#2*^*Pol-γ35*^ and resulted in complete lethality of females and males with either *Act5C-Cas9* or *nos-Cas9*. Note that unlike previously described LOF mutations of *Pol-γ35 (Iyengar et al*., *2002)*, Cas9/gRNA-induced cleavage of the *Pol-γ35* C-terminal domain resulted in embryonic lethality. In fact, we have not observed any larva to emerge from >1000 trans-heterozygous eggs harboring *U6*.*3-gRNA#1*^*Pol-γ35*^ and *nos-* or *Act5C-Cas9*. In sum, these results indicate that both gRNAs are functional and could be used to generate gene drives.

### Development of split HomeR drives with encoded rescue

Using these characterized gRNAs described above, we engineered two *Pol-γ35* HomeR (*HomeR1*^*Pol-γ35*^ and *HomeR2*^*Pol-γ35*^) drives. Fitting with the split-GD design, neither *HomeR1*^*Pol-γ35*^ nor *HomeR2*^*Pol-γ35*^ include the *Cas9* gene, and thus are inherently confineable gene drives. To mediate HDR, left and right homology arm sequences (LHA and RHA) matching the sequences surrounding the cut site of the corresponding gRNA were utilized. Additionally, we included a *3xP3-eGFP-SV40* marker gene to track the presence of the drive and a re-coded C-terminal domain incorporated into the LHA with a p10 3’UTR to support robust expression of the re-coded *Pol-γ35* (**Fig. 2A-C**) and to eliminate homology and prevent gene conversion between the rescue allele and the endogenous allele, which proved problematic in previous drive designs (Champer et al., 2020a, 2020b). Importantly, the recoding was carefully designed to ensure the translation of the re-coded DNA sequence in the wildtype (*wt*) amino acid sequence of *Pol-γ35* with respect to *Drosophila* codon usage bias.

**Figure 2.**
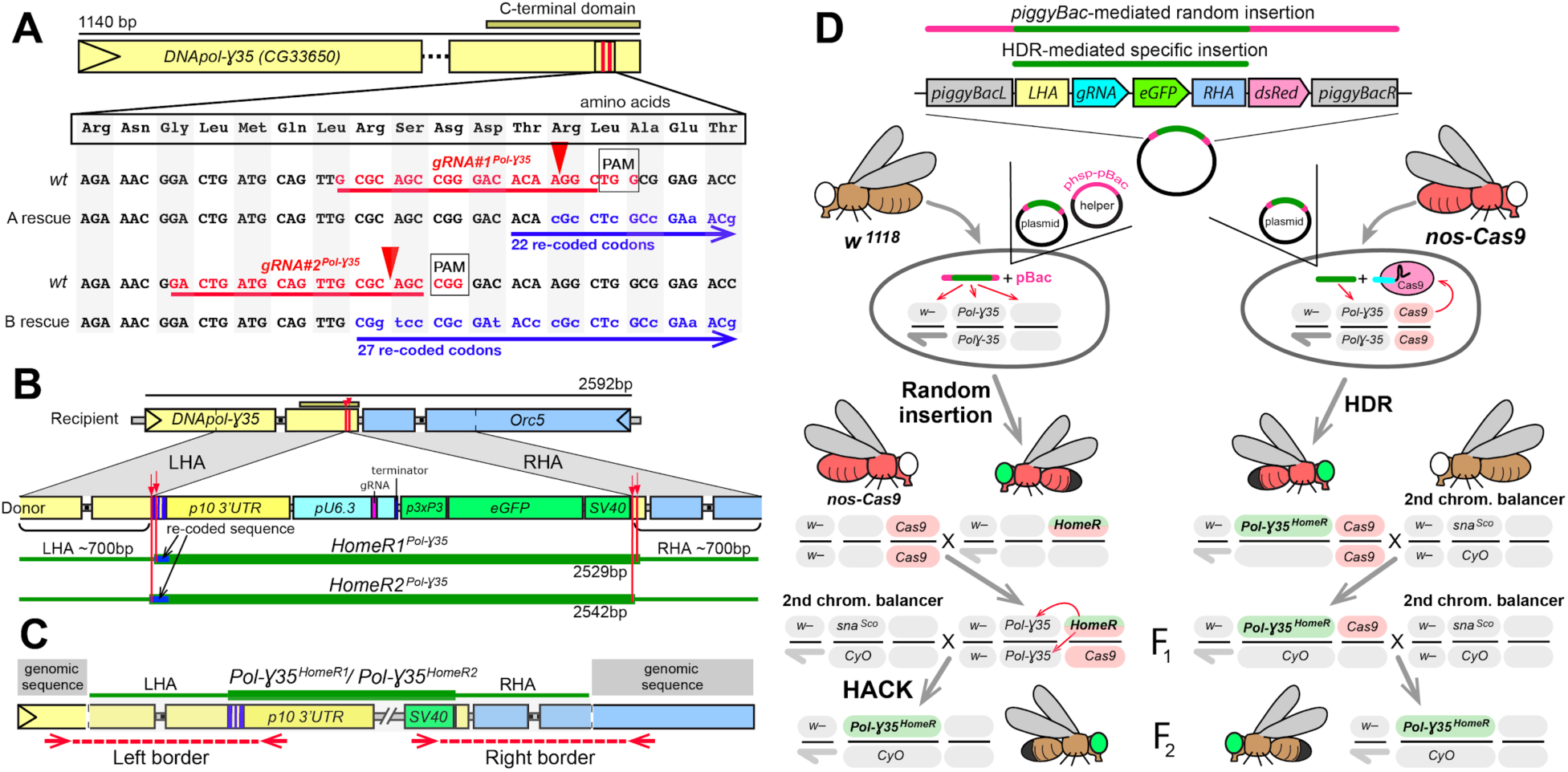
The *Pol-γ35 HomeR* split-drive and its site-specific integration. (**A**) Schematic map of the *DNA polymerase gamma 35 kb gene* (*Pol-γ35*). Two gRNAs targeting its highly conserved C-terminal domain were chosen and tested for guiding Cas9 cleavage (**Fig. 1A-B**). Both gRNA target sites (highlighted in red) are located near the 3’ end of the coding sequence, facilitating re-coding for the rescue allele (highlighted in blue), resistant to Cas9/gRNA-mediated cleavage. Red arrows depict Cas9/gRNA cut sites, which are 13 bases apart. (**B**) Schematic maps of the recipient (wildtype, *wt*) allele encompassing the area spanning *Pol-γ35* and *Orc5* (CG7833) genes, and the donor allele harboring the *HomeR1* or *HomeR2* site-specifically integrated at gRNA cut site #1 or #2 in *Pol-γ35*, respectively. To facilitate site-specific integration, each HomeR genetic construct is surrounded by the Left and Right Homology Arms (LHA and RHA) from the corresponding Cas9/gRNA cut site (red arrows and lines) in the *wt* allele. The re-coded 3’ end sequences of *Pol-γ35* (LHA) are shown in dark blue for both *HomeR1*^*Pol-γ35*^ and *HomeR2*^*Pol-γ35*^. (**C**) To confirm site-specific integration of both *Pol-γ35*^*HomeR1*^ and *Pol-γ35*^*HomeR2*^, the left and right borders between either *HomeR1* or *HomeR2* genetic construct and *Drosophila* genomic sequence around the integration site are sequenced. (**D**) Two separate approaches were used to generate transgenic lines harboring site-specific insertions of *HomeR1* or *HomeR2* at the *Drosophila Pol-γ35*. In the first approach, two plasmids, one carrying the HomeR construct and the helper plasmid (*phsp-pBac (Handler and Harrell, 1999)*) carrying the piggyBac transposase, were injected into *w*^*1118*^ embryos to generate transgenic lines harboring a random piggyBac-mediated integration tagged by double fluorescence (GFP+ and dsRed+). Then transgenic males (GFP+, dsRed+) were crossed to *nos-Cas9* females (dsRed+) to generate site-specific *Pol-γ35*^*HomeR*^ transgenic lines, tagged by GFP alone, using Homology Assisted CRISPR Knock-in (HACK) (Lin and Potter, 2016). In the second approach, the plasmid harboring the HomeR genetic construct was injected alone into *nos-Cas9* embryos (dsRed+) and site-specifically integrated via Homology Directed Repair (HDR) to generate F_1_ heterozygous *Pol-γ35*^*GDe*^/+; nos-Cas9/+ males with double fluorescence (GFP+ and dsRed+). In both approaches, F_1_ transgenic males harboring *Pol-γ35*^*HomeR*^ (GFP+) and *nos-cas9* (dsRed+) were crossed to the 2^nd^ chromosome balancer line, *w*^*1118*^; *CyO/sna*^*Sco*^, to balance and isolate the *Pol-γ35*^*GDe*^ insertion (GFP+).

In case the initial HDR-mediated transgenesis of *HomeR*^*Pol-γ35*^ at the *Pol-γ35* cut site failed, both *HomeR*^*Pol-γ35*^ constructs were assembled in a *piggyBac* plasmid that supported an alternative path for genome integration (**Fig. 2D**). To distinguish between the site-specific HDR-mediated integration, tagged by the eye-specific eGFP fluorescence of *3xP3-eGFP-SV40*, and non-site-specific integration using *piggyBac-*mediated insertion, the *Opie2-dsRed-SV40* marker was included outside the LHA and RHA of both *HomeR*^*Pol-γ35*^, conferring body-specific dsRed fluorescence (**Fig. 2D**). Overall, this split-GD design (Champer et al., 2020a; Kandul et al., 2019a; Li et al., 2020) restricts the spread of *Pol-γ35*^*HomeR*^ and serves as a safeguard against unintended spread, as the homing *Pol-γ35*^*HomeR*^ harboring a *gRNA* and the non-homing *Cas9* are genetically unlinked, resulting in molecular confinement (Esvelt et al., 2014; Marshall and Akbari, 2018).

To generate transgenic lines, we first injected mixtures of each *HomeR*^*Pol-γ35*^ plasmid and a helper plasmid, expressing the pBac transposase that directs random genomic integration (Handler and Harrell, 1999), into *w*^*1118*^ embryos. This established transgenic lines carrying random insertions of the *HomeR1*^*Pol-γ35*^ and *HomeR2*^*Pol-γ35*^ plasmids, genetically tracked by both eGFP and dsRed markers. We then used Homology Assisted CRISPR Knock-in (HACK)(Gantz and Akbari, 2018; Lin and Potter, 2016) to integrate *HomeR1*^*Pol-γ35*^ or *HomeR2*^*Pol-γ35*^ at the corresponding *gRNA*^*Pol-γ35*^ cut site by crossing flies harboring random insertions of *HomeR1*^*Pol-γ35*^ or *HomeR2*^*Pol-γ35*^ to *nos-cas9/nos-Cas9* marked with dsRed (Kandul et al., 2019b) (**Fig. 2D)**. Additionally, to induce HDR-mediated site-specific insertions at the gRNA cut sites in *Pol-γ35* (referred to as *Pol-γ35*^*HomeR*^ once genomically inserted), we injected *HomeR1*^*Pol-γ35*^ or the *HomeR2*^*Pol-γ35*^ plasmid directly into *nos-Cas9/nos-Cas9* embryos (Kandul et al., 2019b) (**Fig. 2D**). One copy of the *Pol-γ35*^*HomeR*^ allele is sufficient to rescue the *wt* function of *Pol-γ35* and, in the presence of Cas9 protein, to support homing in heterozygous *Pol-γ35*^*HomeR*^*/Pol-γ35*^*WT*^ germline cells (**Fig. 2A**).

For both insertion approaches described above (**Fig. 2D**), the F_1_ trans-heterozygous flies harboring potential *Pol-γ35*^*HomeR1*^ or *Pol-γ35*^*HomeR2*^ and tagged by double fluorescence (GFP+ and dsRed+) were individually crossed to *sna*^*Sco*^*/CyO* balancer flies to isolate *Pol-γ35*^*HomeR1*^*/CyO* or *Pol-γ35*^*HomeR2*^*/CyO* flies marked with only GFP fluorescence. Multiple independent transgenic lines of each *Pol-γ35*^*HomeR1*^ and *Pol-γ35*^*HomeR2*^ were isolated and balanced on the second chromosome. To confirm that *Pol-γ35*^*HomeR1*^*/CyO* or *Pol-γ35*^*HomeR2*^*/CyO* lines were indeed inserted at the corresponding cut site in *Pol-γ35*, we assessed their ability for super-Mendelian inheritance in the presence of *Cas9* in *trans* and generated homozygous stocks. Establishment of pure breeding, homozygous stocks of *Pol-γ35*^*HomeR1*^*/Pol-γ35*^*HomeR1*^ and *Pol-γ35*^*HomeR2*^*/Pol-γ35*^*HomeR2*^ demonstrates a functional rescue of *wt Pol-γ35* function. Finally, we sequenced the left and right borders between the *Drosophila* genome and both genetic constructs, the regions spanning the LHA and RHA (**Fig. 2C**), and molecularly confirmed the precision of HDR-mediated insertions at the sequence level.

### Assessment of germline transmission and cleavage rates

To assess the effects of gRNA-mediated cleavage efficiency on transmission rates, we compared the two nearly identical *HomeRs*, as they harbored two distinct gRNA sequences that differed in cleavage efficiencies. The *Pol-γ35*^*HomeR1*^ and *Pol-γ35*^*HomeR2*^ homozygous lines have *U6*.*3-gRNA#1*^*Pol-γ35*^ and *U6*.*3-gRNA#2*^*Pol-γ35*^, respectively, with slightly different LHA and RHA corresponding to their respective gRNA cut sites, which are only 13 bases apart (**Fig. 2A-C**). We found that *Pol-γ35*^*HomeR1*^*/+*; *nos-Cas9/+* trans-heterozygous females crossed to *wt* males transmitted *Pol-γ35*^*HomeR1*^ to 99.5 ± 0.6% of progeny, while *Pol-γ35*^*HomeR2*^*/+*; *nos-Cas9/+* females transmitted the corresponding *Pol-γ35*^*HomeR2*^ to a significantly lower fraction of F_1_ progeny (68.7 ± 6.2%, two-sided Student’s *t*-test with equal variances, *p <* 0.0001; **Fig. 3A, Data S3**). Genetic crosses of either *Pol-γ35*^*HomeR1*^*/+*; *nos-Cas9/+* or *Pol-γ35*^*HomeR2*^*/+*; *nos-Cas9/+* trans-heterozygous males to *wt* females did not result in significant biased transmission to F_1_ progeny (60.7 ± 5.3% vs 52.9 ± 4.0% or 54.3 ± 4.0% vs 51.5 ± 1.8%, respectively, two-sided Student *t*-test with equal variances, *p >* 0.05; **Fig. 3A**). Maternal carryover of Cas9 protein by *nos-Cas9/+* females significantly increased transmission of *Pol-γ35*^*HomeR1*^ by F_1_ *Pol-γ35*^*HomeR1*^*/CyO* females, 66.1 ± 0.8% vs 52.9 ± 4.0% (two-sided Student *t*-test with equal variances, *p <* 0.001; **Fig. 3A, Data S3**). These results suggest that the higher cleavage efficiently of *U6*.*3-gRNA#1*^*Pol-γ35*^, as measured by the induced lethality in the *Lig4* null genetic background (**Fig. 1B**), likely contributes to the higher homing rate of *Pol-γ35*^*HomeR1*^ harboring *U6*.*3-gRNA#1*^*Pol-γ35*^, and underscores the importance of selecting an efficient gRNA for gene drives (GDs).

**Figure 3.**
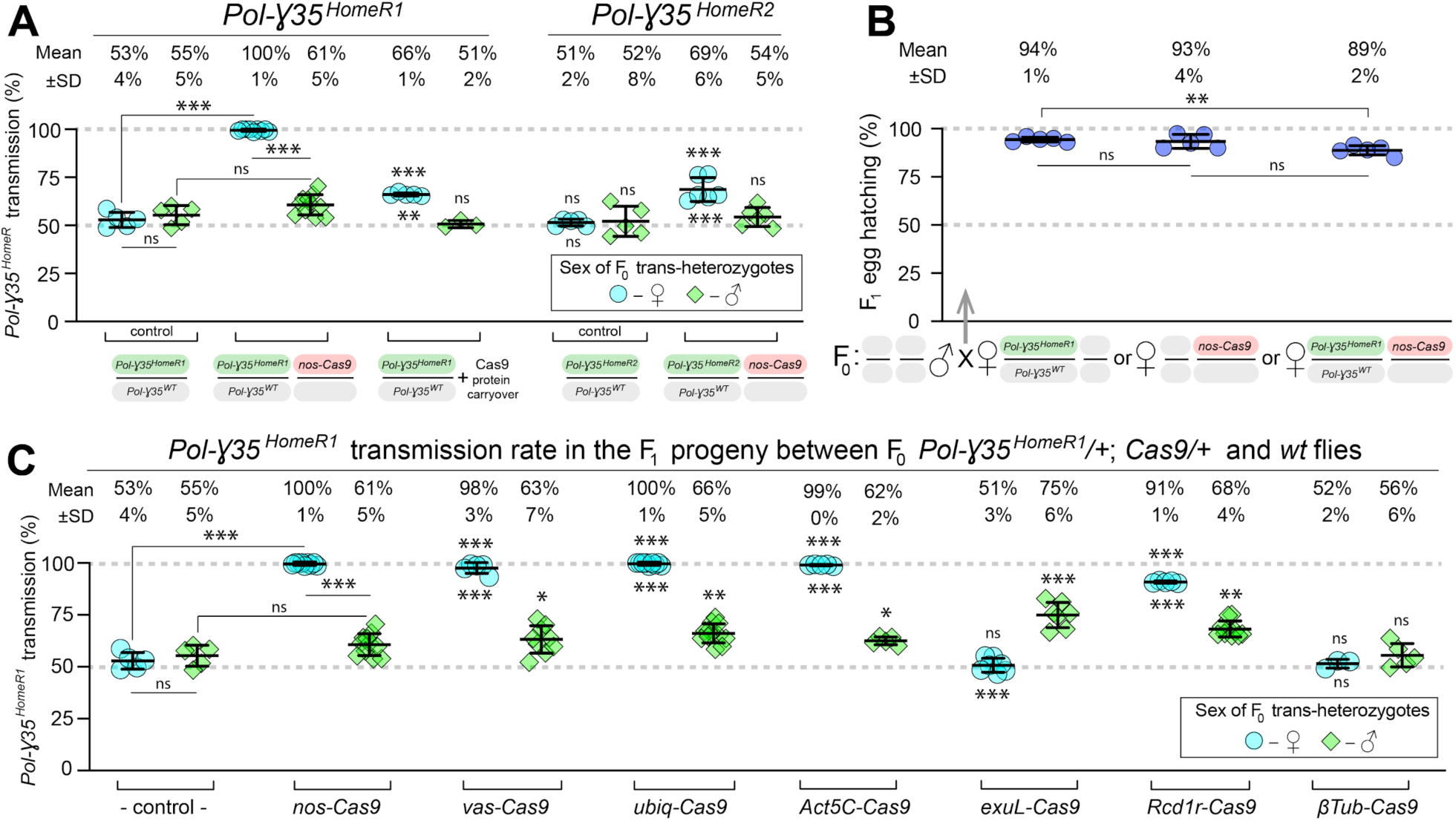
Transmission rates for *Pol-γ35*^*HomeR1*^ and *Pol-γ35*^*HomeR2*^. The *HomeR* element, *Pol-γ35*^*HomeR1*^ or *Pol-γ35*^*HomeR2*^, is inactive by itself and requires Cas9 endonuclease to induce Cas9/gRNA-mediated cleavage for successful homing. This split-drive design permits genetic analysis of a single *HomeR* with different *Cas9* lines. (**A**) Both *Pol-γ35*^*HomeR1*^ and *Pol-γ35*^*HomeR2*^ support super-Mendelian transmission in conjunction with *nos-Cas9* in females, but not in males. *Pol-γ35*^*HomeR1*^ induced significantly higher transmission than *Pol-γ35*^*HomeR2*^, 99.5 ± 0.6% vs 68.7 ± 6.2%, respectively. Notably, maternal carryover of Cas9 protein was sufficient to bias transmission of *Pol-γ35*^*HomeR1*^ by female embryos. (**B**) The hatching rate of F_1_ eggs generated by *Pol-γ35*^*HomeR1*^*/Pol-γ35*^*WT*^; *nos-Cas9/+* females mated to wildtype (*wt*) males was lower by 5% or 4% than that of *Pol-γ35*^*HomeR1*^*/Pol-γ35*^*WT*^; *+/+* or *+/+*; *nos-Cas9/+* females mated to *wt* males (89 ± 2% vs 94 ± 1% or 93 ± 4%, respectively). Therefore, embryonic lethality of *Pol-γ35*^*WT*^ alleles is not the sole mechanism of the nearly ∼100% transmission of *Pol-γ35*^*HomeR1*^. Instead, ∼90% of *wt Pol-γ35* alleles were converted (i.e. homed) into *Pol-γ35*^*HomeR1*^. (**C**) Assessment of different *Cas9* promoters to improve the transmission rate of *Pol-γ35*^*HomeR1*^ in females and males. Trans-heterozygous females (♀) and males (♂) harboring paternal *Cas9* expressed under different promoters were mated to *wt* flies of the opposite sex, and F_1_ progeny were scored for the GFP dominant marker of *Pol-γ35*^*HomeR1*^. The transmission rate was compared to that in *Pol-γ35*^*HomeR1*^*/Pol-γ35*^*WT*^; *+/+* flies without *Cas9* (control) of the corresponding sex (statistical significance indicated above data points). In addition, the transmission rate by trans-heterozygous females was compared to that of trans-heterozygous males for each *Cas9* promoter (statistical significance indicated below data points). Plots show the mean ± SD over at least three biological replicates and rounded to a whole number. Statistical significance was estimated using a two-sided Student’s *t* test with equal variance. (*p* ≥ 0.05^ns^, *p* < 0.05*, *p* < 0.01**, and *p* < 0.001***).

### Majority of *Pol-γ35*^*WT*^ alleles are converted into *Pol-γ35*^*HomeR1*^ alleles in trans-heterozygous females

We hypothesized that either homing (indicating allelic conversion) in oocytes or “destruction” of the *wt* alleles in embryos of trans-heterozygous *Pol-γ35*^*HomeR1*^*/+*; *Cas9/+* females via BLM (Kandul et al., 2019b) could contribute to biased *Pol-γ35*^*HomeR1*^ transmission rates. BLM contributes to RNA-guided dominant biallelic knockouts of the target gene throughout development thereby converting recessive non-functional resistant alleles into dominant deleterious/lethal mutations that can get selected out of a population (**Fig. S1**). Previously, destruction of the *wt* allele in conjunction with maternal carryover of a “toxin” was used to engineer gene drives based on an “addictive” TA approach (Champer et al., 2020a; Oberhofer et al., 2020a, 2019). In these TA drives, one half of the F_1_ progeny did not inherit the TA cassette, meaning not rescued, and were killed—resulting in a rapid spread of the genetic cassette in laboratory populations (**Fig. 6**).

In our experiments, the *U6*.*3-gRNA#1*^*Pol-γ35*^ induced embryonic lethality in the presence of *nos-Cas9* or maternal carryover of the Cas9 protein (**Fig. 1B**). Therefore, to explore the mechanism resulting in the super-Mendelian transmission of *Pol-γ35*^*HomeR1*^, and given that disruption of *Pol-γ35* by *6*.*3-gRNA#1*^*Pol-γ35*^ in *Pol-γ35*^*HomeR1*^ results in fully penetrant embryonic lethality, we determined the egg hatching rate, as the percentage of embryonic lethality, for trans-heterozygous females and compared it to those of females heterozygous for *Pol-γ35*^*HomeR1*^ or *Cas9* (**Fig. 3B**). The hatching rate of F_1_ eggs generated by *Pol-γ35*^*HomeR1*^*/+*; *nos-Cas9/+* trans-heterozygous females crossed to *wt* males was reduced by 5% as compared to *Pol-γ35*^*HomeR1*^*/+*; *+/+* heterozygous females crossed to *wt* males (88.8 ± 2.4% vs 94.4 ± 1.2%; two-sided Student *t*-test with equal variances, *p <* 0.004). Furthermore, the hatching rate of eggs laid by trans-heterozygous females was not statistically different from that laid by *+/+*; *nos-Cas9/+* heterozygous females crossed to *wt* males (88.8 ± 2.4% vs 93.4 ± 3.7%; two-sided Student’s *t*-test with equal variances, *p <* 0.052, **Fig. 3B, Data S4**). Moreover, there was no significant difference between the larvae-adult survival rates comparing *Pol-γ35*^*HomeR1*^/+; *nos-Cas9*/+ to *Pol-γ35*^*HomeR1*^/+; +/+ indicating that there is no bias at these later stages. Taken together, these data indicate that from the expected 50% of *Pol-γ35*^*WT*^ alleles transmitted by trans-heterozygous females, ∼5% were “destroyed” via BLM—meaning mutated and not complemented by the paternal allele, since it was also mutated by Cas9/gRNA maternal carryover (**Fig. S2**)—and the remaining ∼45% were converted into *Pol-γ35*^*HomeR1*^—resulting in an estimated conversion rate of∼90% (45% X 2). Therefore, these data indicate that the observed transmission rate of nearly ∼100% was caused by ∼90% conversion and ∼10% “destruction” of the *Pol-γ35*^*WT*^ alleles. In sum, the *Pol-γ35*^*HomeR1*^ transmission rate of nearly ∼100% observed in *Pol-γ35*^*HomeR1*^*/+*; *nos-Cas9/+* trans-heterozygous females could not be simply explained by the “destruction” of all *wt Pol-γ35* alleles, which would result in the lethality of 50% progeny as in non-homing ClvR (Oberhofer et al., 2020a, 2020b, 2019) and TARE (Champer et al., 2020a) drives, and instead is a result of both conversion and destruction of the recipient allele at the *Pol-γ35* locus (**Fig. 2B**).

### *Nos-* and *ubiq-Cas9* support the strongest female-specific transmission of *Pol-γ35*^*HomeR1*^

The split-drive design facilitates testing of different Cas9 promoters. Therefore, we were able to estimate the transmission of *Pol-γ35*^*HomeR1*^ by trans-heterozygous females and males harboring *Pol-γ35*^*HomeR1*^ in combination with four alternative Cas9 promoters active in germ cells of both sexes. *Nanos* (*nos*) and *vasa* (*vas*) promoters were previously described as germline-specific promoters active in both sexes (Hay et al., 1988; Sano et al., 2002; Van Doren et al., 1998), though recent evidence indicates ectopic expression in somatic tissues from both *nos-Cas9* and *vas-Cas9* (Kandul et al., 2019a, 2019b). The *Ubiquitin 63E (Ubiq)* and *Actic 5C* (*Act5C)* promoters in *ubiq-Cas9 (Kandul et al*., *2019b)* and *Act5C-Cas9 (Port et al*., *2014)* transgenic lines, respectively, support strong expression in both somatic and germ cells (Kandul et al., 2019a, 2019b; Port et al., 2014; Preston et al., 2006). To control for genome insertion effects, each *Cas9* transgene was inserted at the same attP docking site on the 3rd chromosome, except for *Act5C-Cas9* that was integrated on the X chromosome (Port et al., 2014). Since maternal carryover of the Cas9 protein was shown to induce a “shadow drive” two generations later (Guichard et al., 2019; Kandul et al., 2019a), we used trans-heterozygous flies that inherited paternal *Cas9* to quantify the transmission of *Pol-γ35*^*HomeR1*^. Trans-heterozygous females carrying *Pol-γ35*^*HomeR1*^ together with *nos-Cas9, vas-Cas9, ubiq-Cas9*, or *Act5C-Cas9* crossed to *wt* males biased transmission of *Pol-γ35*^*HomeR1*^ to nearly ∼100% of F_1_ progeny (99.5 ± 0.6%, 97.6 ± 2.6%, 99.6 ± 0.6%, and 99.0 ± 0.4%, respectively, vs 52.9 ± 4.0% by *Pol-γ35*^*HomeR1*^*/Pol-γ35*^*WT*^; *+/+* females, two-sided Student’s *t*-test with equal variances, *p <* 0.001; **Fig. 3C**). Note that the corresponding trans-heterozygous males only modestly biased *Pol-γ35*^*HomeR1*^ transmission from 55.3 ± 5.0% of F_1_ progeny to 60.7 ± 5.3% (*p* > 0.05), 63.2 ± 6.6% (*p* < 0.03), 66.1 ± 4.6% (*p* < 0.004), and 62.0 ± 1.7% (*p* < 0.017, two-sided Student’s *t*-test with equal variances, **Fig. 3C, Data S3**), respectively.

### *ExuL-Cas9* supports the strongest male-specific transmission of *Pol-γ35*^*HomeR1*^

To assess whether males could support robust homing similar to females, we investigated three alternative male-specific promoters. We established the *Drosophila exuperantia* (CG8994) large fragment (*exuL*) promoter for an early male-specific expression. The *Rcd-1 related* (*Rcd1r*, CG9573)(Chan et al., 2013) and *βTubulin 85D (βTub)(Chan et al*., *2011; Michiels et al*., *1989)* promoters support an early and late, respectively, testis-specific expression in *Drosophila* males. We found that only *exuL-Cas9* induced the male-specific super-Mendelian inheritance of *Pol-γ35*^*HomeR1*^; trans-heterozygous males, but not females, transmitted *Pol-γ35*^*HomeR1*^ to more than 50% of F_1_ progeny (75.0 ± 6.1% vs 55.3 ± 5.0% in ♂, *p <* 0.0001; and 50.7 ± 3.4 % vs 52.8 ± 4.0% in ♀, *p >* 0.05, two-sided Student’s *t*-test with equal variances; **Fig. 3C, Data S3**). To our surprise, *Rcd1r-Cas9* induced super-Mendelian inheritance of *Pol-γ35*^*HomeR1*^ in both trans-heterozygous males and females (68.2 ± 3.8% vs 55.3 ± 5.0% in ♂, *p <* 0.002; and 90.8 ± 0.5% vs 52.8 ± 4.0% in ♀, *p >* 0.0001, two-sided Student’s *t*-test with equal variances; **Fig. 3B**). Finally, *βTub-Cas9* did not induce changes in transmission of *Pol-γ35*^*HomeR1*^ by either trans-heterozygous males or females (55.6 ± 5.7% vs 55.3 ± 5.0% in ♂, *p =* 0.55; and 51.5 ± 2.1% vs 52.9 ± 4.0% in ♀, *p =* 0.94, two-sided Student’s *t*-test with equal variances; **Fig. 3B, Data S3**). These results suggest that *Drosophila* males bias *Pol-γ35*^*HomeR1*^ transmission, however this bias is substantially lower than the nearly ∼100% transmission of *Pol-γ35*^*HomeR1*^ in females.

### Functional *Pol-γ35* resistance alleles (*Pol-γ35*^*R1*^) did not hinder drive persistence in 10 generations

We reasoned that insertion of *HomeR* into the gene required for viability could also prevent the accumulation of *Pol-γ35* LOF resistance alleles (R2 type, *Pol-γ35*^*R2*^) by exploiting BLM (**Fig. S1**). However, functional resistance alleles (R1 type, *Pol-γ35*^*R1*^), either from in-frame *indels* or synonymous base substitutions (SBS), could still be induced by *Cas9/gRNA#1*^*Pol-γ35*^. This could be problematic if *Pol-γ35*^*R1*^ resistance alleles do not impose fitness costs on homozygous carriers, as they would be expected to spread at the expense of the drive. To explore this potential, we set up three laboratory populations of heterozygous *Pol-γ35*^*HomeR1*^*/+* flies in the *nos-Cas9/nos-Cas9* genetic background and assessed the emergence and spread of induced resistance alleles over ten discrete generations (**Fig. 4A**). Although we could not distinguish homozygous flies from heterozygotes with respect to the dominant marker of *Pol-γ35*^*HomeR1*^, functional and fit *Pol-γ35*^*R1*^ alleles were expected to spread at the expense of *Pol-γ35*^*HomeR1*^ alleles, which would be straightforward to score in our assay by loss of the GFP marker and would be predicted to become homozygous over multiple generations.

**Figure 4.**
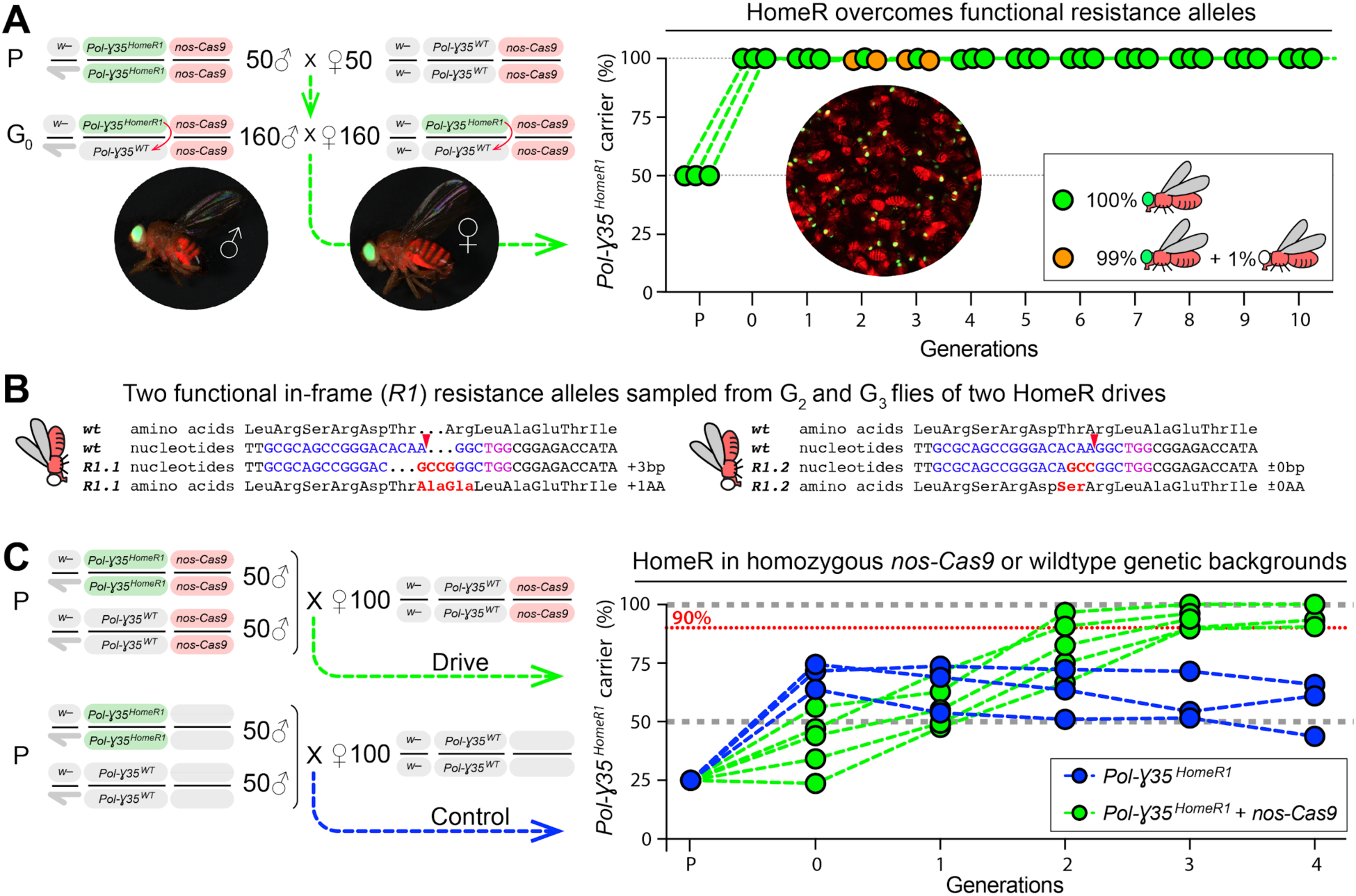
Functional resistance alleles (*Pol-γ35*^*R1*^) do not impede the spread of *Pol-γ35*^*HomeR1*^. (**A**) To explore the fate of induced functional resistance alleles (*R1*), three gene population cage experiments were set up starting with *Pol-γ35*^*HomeR1*^*/Pol-γ35*^*WT*^ heterozygous flies in *nos-Cas9/nos-Cas9* genetic background and run for ten discrete generations. *Pol-γ35*^*HomeR1*^*/Pol-γ35*^*WT*^; *nos-Cas9/nos-Cas9* flies were tagged by eye-specific GFP and body-specific dsRed. Images of an individual male (♂), female (♀), and a group of flies are shown. In total, nine viable flies (lacking the *Pol-γ35*^*HomeR1*^ allele), as determined by the absence of dominant eye-specific GFP expression, were identified at generations 2 and 3 in cages #1 and #3. After these flies were allowed to mate and lay eggs for the next generation, then isolated and genotyped. (**B**) None of nine genotyped flies harbored the *Pol-γ35*^*WT*^ allele. Instead, each fly carried at least one functional resistance allele rescuing the viability of these flies. Both different types of *R1* alleles change the amino acid sequence. Seven of nine flies were heterozygous, harboring one of the identified *R1* alleles together with an out-of-frame *indel* allele. The sequence of *gRNA#1*^*Pol-γ35*^ is highlighted in blue, and its PAM sequence is in purple. Red arrows depict Cas9/gRNA cut sites. Base insertions and amino acid changes are in red. (**C**) To assess the performance of *HomeR* gene drive, population drive lineages were seeded with 50 *Pol-γ35*^*HomeR1*^*/Pol-γ35*^*HomeR1*^ ♂, 50 *wt* ♂, and 100 *wt* virgin ♀ in the presence (green points) or absence (blue points) of *nos-Cas9*, and the carrier frequency of *Pol-γ35*^*HomeR1*^ was scored by a dominant 3xP3-GFP marker each discrete generation. In the span of four generations of drive, *Pol-γ35*^*HomeR1*^ allele spread from the introductory frequency of 25% to the carrier frequency of 94.8 ± 3.5% in the presence of *nos-Cas* (green points and dotted lines) or 56.9 ± 11.6% without the *Cas9* transgene (*p* = 0.025, a two-sided Student’s *t* test with unequal variance). The model for the *HomeR* population replacement drive (gray points and a black dotted line) was fitted to the empirical data of the *Pol-y35^HomeR1^* spread in the presence of *nos-Cas9* (green points).

Out of 10 generations with three distinct populations, we found only nine flies lacking the *Pol-γ35*^*HomeR1*^ allele (i.e. GFP negative) at generations 2 and 3 in two out of three population lineages (drives #1 and #3, **Fig. 4A; Data S5**). To rescue their viability in the absence of *Pol-γ35*^*HomeR1*^, these flies had to harbor either *Pol-γ35*^*WT*^ or *Pol-γ35*^*R1*^ and were analyzed to determine their genotype. To ensure that any generated *Pol-γ35*^*R1*^ alleles had a chance to spread and compete with *Pol-γ35*^*HomeR1*^ alleles, these flies were transferred among the subsequent generation and allowed to mate with other flies and lay eggs in each population lineage before they were genetically analyzed. The analysis revealed that each fly harbored at least one *Pol-γ35*^*R1*^ resistance allele that rescued viability. Two different *Pol-γ35*^*R1*^ alleles were identified by sequencing multiple clones of amplicons from each of the nine genotyped flies. One *Pol-γ35*^*R1*^ allele was sampled in eight flies from two independent lineages. It had a three-base-insertion that inserted one amino acid as well as a change in one amino acid, p.T358_L360insAG (R1.1, **Fig. 4B**). The other *Pol-γ35*^*R1*^ allele was found in two flies, and its three-base-substitution caused one amino acid change, p.T358S (R1.2, **Fig. 4B**). Seven out of nine genotyped flies were heterozygous, harboring both functional and LOF alleles at the *Pol-γ35* locus, one fly had two different *Pol-γ35*^*R1*^ alleles, and one fly could be homozygous for the *R1*.*1* allele, since ten sequenced clones gave the same *R1*.*1* allele (**Fig. 4B**). We did not identify any fly without the *Pol-γ35*^*HomeR1*^ allele after generation 3, as indicated by the eye-specific GFP expression (**Fig. 4A, Data S5**), thus the identified *Pol-γ35*^*R1*^ alleles did not spread and we were unable to establish these as isolated strains, indicating that flies harboring these alleles were likely less competitive than those with one copy of the *Pol-γ35*^*HomeR1*^ allele as would be expected when targeting a haplosufficient gene.

To further explore the diversity of resistance alleles remaining after ten generations we performed next-generation sequencing on sixty randomly chosen flies (each fly had at least one copy of *Pol-γ35*^*HomeR1*^) from each drive to identify and quantify any “short” *Pol-γ35* alleles, which did not harbor the large insert of *HomeR1*^*Pol-γ35*^ (∼2.5 KB, **Fig. 2A**). Neither the *Pol-γ35*^*WT*^ nor previously identified *Pol-γ35*^*R1*^ alleles were sampled among nearly 150,000 sequence reads for the three drive experiments. Instead, we found two novel in-frame *indel* alleles, 18 bp and 9 bp deletions, in drives #2 and #3 (**Fig. S2**). The 18 bp deletion was responsible for 80% of the resistance alleles sampled for drive #2, while the 9 bp deletion was the least abundant (5%, **Fig. S2**) resistance allele in drive #3. Since both in-frame *indel* alleles cause the deletion of either 6 or 3 amino acids in the middle of the highly conserved c-terminal domain of *Pol-γ35*, they are likely deleterious recessives. The remaining eleven alleles were out-of-frame *indels*, ranging from a 1 bp insertion to a 23 bp deletion (**Fig. S2**). Two LOF alleles, 2 bp and 4 bp deletions, were also seen in the genotyped flies at generations 2 and 3. The relative abundance of each allele can be used to extrapolate the minimum number of resistance alleles sampled in the sixty heterozygous and/or homozygous flies and thus persisted though ten generations of drive. We inferred that at least 9, 5, and 17 resistance alleles persisted for ten generations and were rescued by the *Pol-γ35*^*HomeR1*^ allele in 60 sampled flies from drives #1, #2, and #3, respectively (**Fig. S2**). The fact that these induced functional resistance alleles did not take over at the expense of *HomeR1* in 10 generations (i.e. the drives were all at genotypic fixation) indicates that these alleles could not compete with the *Pol-γ35*^*HomeR1*^ re-coded rescue and spread at its expense.

### *HomeR* spreads efficiently into small populations

The *Pol-γ35*^*HomeR1*^ drive spreads in experimental populations in the presence of *Cas9*. To evaluate the drive efficiency of *HomeR*, we established five drive and three control (‘no-drive’) populations by seeding 50 homozygous *Pol-γ35*^*HomeR1*^ males and 50 *wt* males together with 100 *wt* virgin females in the presence (homozygous *nos-Cas9*) or absence of *Cas9* (**Fig. 4C**). The introduction ratio of *Pol-γ35*^*HomeR1*^ to *Pol-γ35*^*WT*^ was 25% (1:3) in the parental generation (P). Note that released males were competing for mating with virgin females, and their mating competitiveness could be scored by the dominant 3xP3-GFP marker of *Pol-γ35*^*HomeR1*^ in their progeny at generation 0 (G_0_). Both types of homozygous *Pol-γ35*^*HomeR1*^ males with and without *Cas9* were able to compete with the corresponding *wt* males indicating that the reconded part of *Pol-γ35* and an exogenous p10 3’UTR rescued the *wt* function of Pol-γ35 without causing a strong fitness cost (**Fig. 4C**). The *Pol-γ35*^*HomeR1*^ drive has spread to 100% carrier frequency in one out of five drive populations by generation 3, and climbed above 90% in the remaining four drive populations by generation 4 (**Fig. 4C**). In the absence of *Cas9*, the *Pol-γ35*^*HomeR1*^ drive has spread to the significantly lower frequency in three control populations (56.9 ± 11.6% in non-drive vs 94.8 ± 3.5% in drive population at generation 4, *p* = 0.025, a two-sided Student’s *t* test with unequal variance), indicating Cas9 dependence for drive.

Fitting a mathematical model of CRISPR/Cas9-based homing drive to the observed cage data (see Methods), we found the data to be consistent with cleavage efficiencies in females and males of 98.8% (95% credible interval (CrI): 95.2-99.9%) and 99.3% (95% CrI: 97.2-100%), respectively, and a frequency of accurate HDR given cleavage in females and males of 99.3% (95% CrI: 97.0-100%) and 9.3% (95% CrI: 7.2-10.0%), respectively. When accurate HDR did not occur, the data were consistent with 1.1% (95% CrI: 0.1-4.2%) of resistant alleles being in-frame, and the remainder being out-of-frame or otherwise costly LOF alleles. Individuals having the *HomeR* system were found to have a negligible fitness cost of 0.4% (95% CrI: 0.0-2.1%), while individuals homozygous for the LOF allele were modeled as completely unviable. The fitted parameter estimates are consistent with parameters directly estimated in this study, and the fitted model trajectory of GFP+ individuals is consistent with the observed cage data depicted in **Fig. 4C**.

### Modeling indicates that *HomeR* is an efficacious gene drive

To compare the performance of HomeR against contemporary gene drive systems for population modification, we modeled one-(aka. autonomous) and two-locus (aka. split-drive) versions of ClvR (Oberhofer et al., 2020a, 2020b, 2019), the one-locus TARE system from (Champer et al., 2020a) as well as a two-locus TARE configuration based on their design, a HGD targeting a non-essential gene (Gantz et al., 2015; Hammond et al., 2016), and HomeR. In each case, we first simulated population spread of each gene drive system for an ideal parameterization (see Methods for more details) and included additional simulations for HomeR under current experimentally-derived parameters (HomeR-exp, **Fig. 5A and 5C**). To gauge behavior across a range of scenarios, we performed simulations for a range of fitness costs (implemented as female fecundity reduction) and drive system transmission rates (implemented by varying the cleavage rate), providing heatmaps of the expected performance for each drive system at each parameter combination (**Fig. 5B and 5D**). Drive efficacy, the outcome in these comparisons, is defined as the expected fraction of individuals that carry the drive (and a linked effector) allele, in either heterozygous or homozygous form, 20 generations following a 25% release of female and male heterozygotes for each drive system.

**Figure 5.**
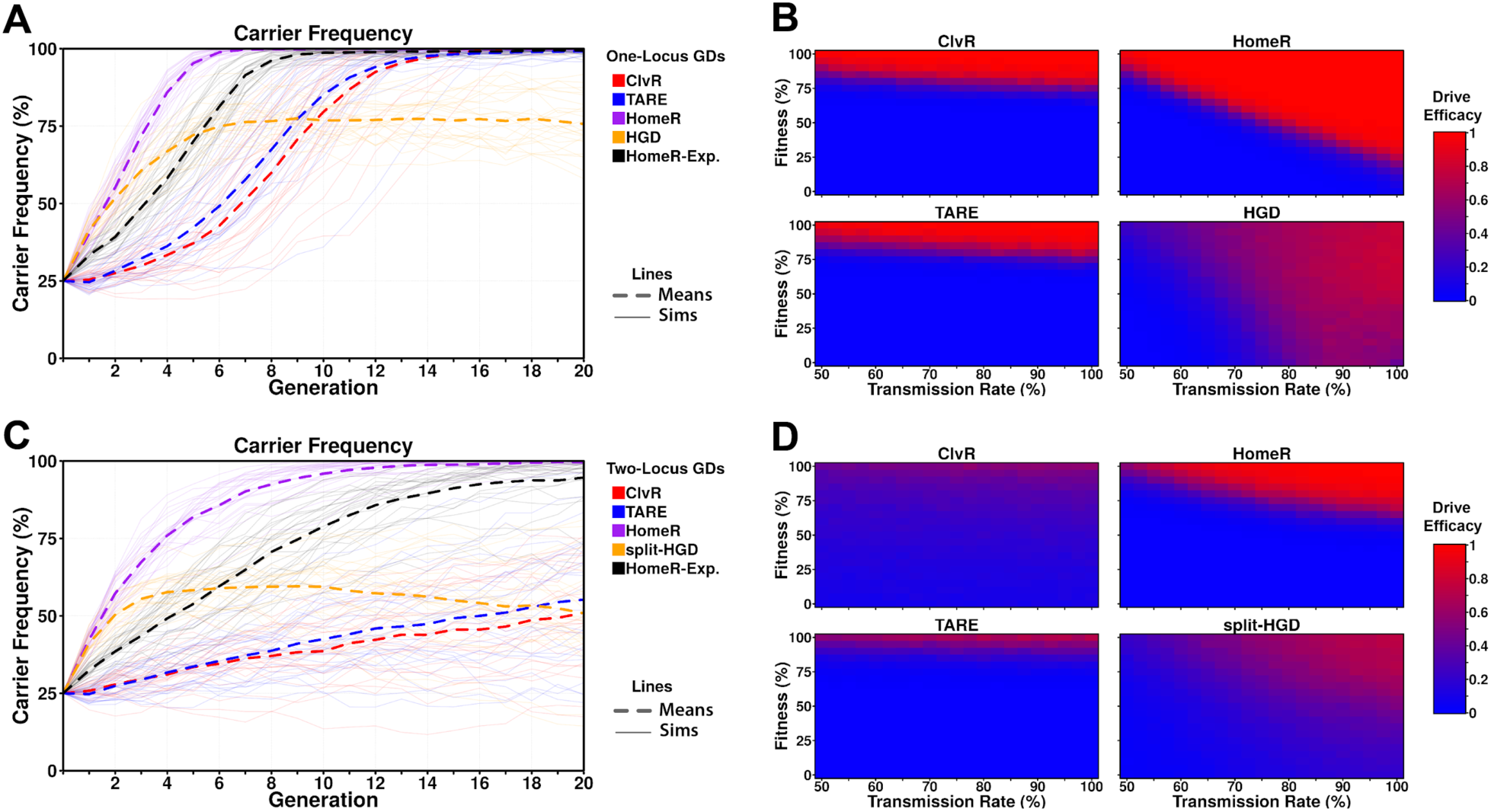
Performance of contemporary gene drive systems for population modification. (**A**) Simulations of carrier frequency trajectories (i.e. heterozygotes and homozygotes) for one-locus versions of ClvR, TARE, HomeR, and HGD for ideal parameters (see Methods), and HomeR for experimental parameters (HomeR-Exp, see Methods). 25 repetitions (lighter lines) were used to calculate the average behavior of each drive (thicker, dashed lines). Populations were initialized with 75% wildtype (*+/+*) adults and 25% drive heterozygotes (*drive/+*), equally split between females and males. (**B**) Heatmaps depicting drive efficacy for one-locus versions of ClvR, TARE, HomeR, and HGD for a range of fitness and transmission rate parameter values. Fitness costs were incorporated as a dominant, female-specific fecundity reduction. Transmission rate was varied based on cleavage rate, using HDR rates consistent with ideal parameters, when applicable (see Methods). Drive efficacy is defined as the average carrier frequency at generation 20 (approximately 1 year, given a generation period of two to three weeks) based on 100 stochastic simulations with the same initial conditions as **A**. (**C**) Simulations of carrier frequency trajectories for two-locus (split-drive) versions of ClvR, TARE, HomeR, and HGD for ideal parameters (see Methods), and HomeR for experimental parameters (HomeR-Exp, see Methods). 25 repetitions (lighter lines) were used to calculate the average behavior of each drive (thicker, dashed lines). Populations were initialized with 75% wildtype (*+/+; +/+*) adults and 25% drive heterozygotes (*Cas9/Cas9; gRNA/+*), equally split between females and males. (**D**) Heatmaps depicting drive efficacy for two-locus versions of ClvR, TARE, HomeR, and HGD for a range of fitness and transmission rate parameter values, implemented as in panel **B**. Drive efficacy is defined as the average carrier frequency at generation 20 (approximately 1 year, given a generation period of two to three weeks) based on 100 stochastic simulations with the same initial conditions as **C**.

When one-locus GD systems are compared for ideal parameter values, HomeR outperforms all other GDs in terms of speed of spread, and reaches near fixation in terms of carrier frequency, as does ClvR and TARE (**Fig. 5A**). HGD displays a similar speed of spread to HomeR initially; however, fitness costs from the targeted gene knockout and loss-of-function (R2) alleles build up over time and progressively reduce its speed of spread and efficacy. The HomeR design overcomes this fitness reduction and R2 allele buildup by rescuing the *wt* function of a targeted essential gene (**Fig. 6**). ClvR and TARE perform similarly to each other for ideal parameter values, but reach near carrier fixation ∼8-9 generations after HomeR does for ideal parameter values (**Fig. 5A**). When experimental parameters are used for HomeR (HomeR-Exp, in **Fig. 5A**), it reaches near carrier fixation ∼3 generations later than for ideal parameter values; but still spreads faster than the ClvR and TARE systems with ideal parameters. HomeR also reaches near carrier fixation for the widest range of fitness and transmission rate parameter values (**Fig. 5B**), outperforming all alternative drives by this criterion. As a HDR, HomeR drives to high carrier frequencies provided its inheritance bias (or transmission rate) exceeds its associated fitness cost. Drive efficacy of ClvR and TARE, on the other hand, is strongly dependent on fitness cost and weakly dependent on transmission rate. Indeed, ClvR and TARE can each only tolerate fitness costs less than ∼20% (**Fig. 5B**). This is a consequence of their design, employing a toxin-antidote scheme, which induces a significant fecundity reduction (**Fig. 6A-B**) in addition to other fitness costs. A one-locus HGD also exhibits efficacy across a wide range of parameter combinations, but its efficacy is reduced compared to HomeR due to the build-up of R2 and R1 alleles (**Fig. 6C-D**), which can potentially block spread of the HGD in large populations (**Fig. 5B**).

**Figure 6.**
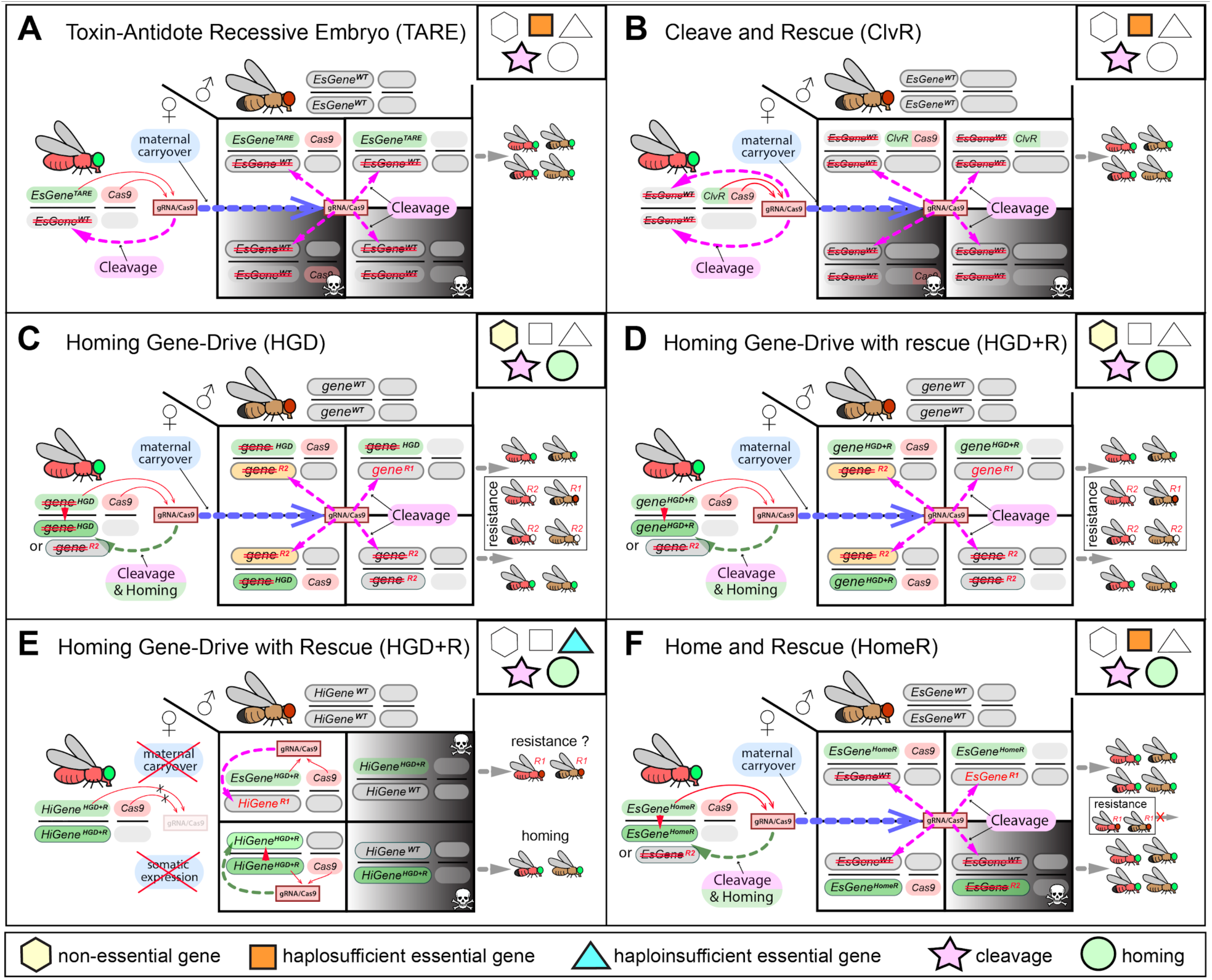
Mechanistic comparison of contemporary split-drives for population modification. Each diagram depicts the cross between females trans-heterozygous for a GD and wildtype (*wt*) males. <u>T</u>oxin-<u>A</u>ntidote <u>R</u>ecessive <u>E</u>mbryo (TARE) (**A**) and <u>Cl</u>ea<u>v</u>e and <u>R</u>escue (ClvR) (**B**) are non-homing TA-based drives. TARE and ClvR force their inheritance by knocking out an essential gene (*EsGene*^*WT*^) in oocytes as well as in embryos by the maternal carryover of Cas9/gRNA, and rescuing only those embryos that inherit TARE and ClvR genetic cassettes harboring a re-coded essential gene, which is resistant to Cas9/gRNA-mediated cleavage. As a result, mating between trans-heterozygous TARE and ClvR females and *wt* males generate 50% non-viable embryos. The TARE is integrated at the target gene locus and uses its native promoter to drive expression of the re-coded rescue, hence it is referred to as *EsGene*^*TARE*^. Panel **A** shows its two-locus version, in which Cas9 is expressed from a separate chromosome. Both components of a two-locus ClvR are inserted at genomic loci separate from the target gene. Panel **B** depicts the two-locus version of ClvR, in which a ClvR harbors a re-coded rescue with its sequence-distinct promoter and 3’UTR, and both ClvR and Cas9 are inserted into two distant loci. Since both TARE and ClvR use multiple gRNAs to target an essential gene, only very rare functional resistant alleles (R1) can survive. (**C**) A homing gene-drive (HGD) spreads its inheritance in heterozygous germ cells by cleaving a non-essential gene (*gene*) and homing, or copying itself, at the cut site (*gene*^*GD*^). Since the knockout of a non-essential gene does not cause lethality and sterility of *gene*^*GD*^*/gene*^*GD*^, HGD spreads itself, though the fitness of *gene*^*GD*^*/gene*^*GD*^ is lower than that of *gene*^*WT*^*/gene*^*GD*^ or *gene*^*WT*^*/gene*^*WT*^. Maternal carryover of Cas9/gRNA knocks out paternal alleles in embryos, and NHEJ induces large levels of both R2 and R1 (*gene*^*R2*^ and *gene*^*R1*^) resistance alleles that survive and eventually block the spread of HGD. (**D**) The HGD with a rescue (HGD+R) preserves the *wt* function of a knocked out non-essential gene after its precise insertion via homing and improves the spread, however incomplete sterility of homozygous loss-of-function (R2) alleles results in accumulation of resistance alleles impeding the spread of HGD+R. (**E**) The HGD+R inserted at a haploinsufficient gene (*HiGene*) requires both alleles to express functional transcripts for viability and fertility of its carriers. Any cleavage that does not result in precise homing very early during the development of trans-heterozygous embryos *HiGene*^*HGD+R*^*/HiGene*^*WT*^; *Cas9/+* will induce high fitness cost or lethality via lethal mosaicism. Therefore, the maternal carryover and somatic expression of *Cas9/gRNA* complexes, which are empirically unavoidable, make engineering and proper functionality of *HiGene*^*HGD+R*^ unachievable. (**F**) <u>Home</u> and <u>R</u>escue (HomeR) drive as a toxin-antidote homing drive. HomeR harbors a re-coded essential gene, and its precise homing at the cut site rescues the *wt* function of the essential gene (*EsGene*^*HomeR*^). Maternal carryover of HomeR’s Cas9/gRNA induces cleavage of paternal *EsGene*^*WT*^ alleles in embryos, that are rescued by only *EsGene*^*HomeR*^ but not *EsGene*^*R2*^ maternal alleles resulting in the removal of non-rescued loss-of-function NHEJ alleles (*EsGene*^*R2*^) via biallelic lethal mosaicism (See **Fig. S1**). HomeR targets an ultra-conserved region of an essential gene for knockout to minimize induction of functional resistance alleles (*EsGene*^*R1*^) that are not fitness costly. Since HomeR is less costly to the female fertility than TARE and ClvR, and induces less resistance alleles that a homing GD, HomeR outperforms contemporary gene drives (**Fig. 5**). Red strikethrough defines a LOF (R2) allele.

In two-locus simulations, Cas9 is separated from the gRNAs in all designs and undergoes independent assortment during gametogenesis. The effects of this design change are evident (**Fig. 5C**). Under the same experimental conditions as one-locus simulations, there is significantly more variation in behavior of two-locus GDs, with a reduced speed of introgression into the population and slightly reduced overall efficacy. Nevertheless, HomeR still demonstrates strong performance, spreading significantly faster than ClvR and TARE. TARE performs significantly worse in a split configuration (Champer et al., 2020a). ClvR, when completely unlinked, also performs significantly worse, in-agreement with results from (Oberhofer et al., 2020a). Exploring the performance under a range of parameters, we found reduced efficacy for all drives (**Fig. 5D**); however, trends of between-drive performance are maintained in this new configuration. TARE and ClvR drive efficacy is still nearly independent of transmission rate, but sharply dependent on fitness costs, rarely reaching carrier fixation for the explored parameter values. HGD still exhibits efficacy across a wide range of parameter combinations, but efficacy is limited in all of them. HomeR still demonstrates higher efficacy than other drives, but for a smaller range of parameter combinations.

## Discussion

We have engineered a system we term HomeR, for population modification that mitigates existing issues related to drive resistance. To limit the potential for inducing functional resistance alleles, an ultraconserved, haplosufficient gene required for insect viability was targeted. Multigenerational population drive experiments indicate that *Pol-γ35*^*HomeR*^ can spread and persist efficiently in the presence of Cas9, and this persistence is not impacted by induced resistance alleles, including functional resistance alleles, overcoming a major challenge for population modification HGDs.

The re-coded rescue strategy that we used to develop HomeR was also used in previous *Drosophila* toxin-antidote non-homing GDs (Champer et al., 2020a; Oberhofer et al., 2020a, 2020b, 2019) and recent HGD’s in both *Drosophila* (Champer et al., 2020b), and *Anopheles stephensi (Adolfi et al*., *2020)*, though each of these examples suffered from potential drawbacks. For example, both the haplolethal HGD (Champer et al., 2020b) and the TARE design (Champer et al., 2020a) share similar problematic design architectures that can be unstable as they are susceptible to functional resistance alleles induced via recombination between the promoter including sequences 5’ of the coding sequence and 3’UTR regions, which are identical between the re-coded sequence and the *wt* sequence (Fig. S2C in (Champer et al., 2020a) and Fig. 2 (Champer et al., 2020b)). Moreover, the haplolethal HGD (Champer et al., 2020b) requires a strict germline specific promoter that lacks maternal carryover (**Fig. 6E**) otherwise lethal mosaicism (**Fig.1C, Fig. S1**), either mono- or bi-allelic, will result in dominant negative fitness costs to its carrier and impede drive spread or inheritance. In fact, our efforts to find such promoters in *Drosophila* proved exceedingly difficult - with previously tested “germline specific” promoters such as *nanos* and *vasa* showing significant somatic activity at multiple insertion sites (Kandul et al., 2019a, 2019b). The recent HGD in *Anopheles stephensi (Adolfi et al*., *2020)* was designed to target and rescue a non-essential gene for viability (i.e. the eye pigmentation *kynurenine hydroxylase* (*kh*) gene), whose knockout was pleiotropic and only partially costly to female fecundity and survival (Adolfi et al., 2020; Gantz et al., 2015; Pham et al., 2019). Notwithstanding, this drive spread efficiently in small, multigenerational laboratory population cages under several release thresholds, however, many drives did not reach, nor maintain, complete fixation presumably due to the viability and partial fertility of drive generated homozygous LOF resistance alleles (**Fig. 6D**), underscoring the critical importance of targeting a recessive haplosufficient essential gene for such drives especially for larger releases. Comparatively, the ClvR system is quite stable, however it can be cumbersome to engineer —requiring re-coding of the essential rescue gene, including all target sequences within the coding sequence (lacking introns), and uses an exogenous promoter and 3’UTR, necessitating precise titration of expression from a distal genomic location with exogenous sequences to guarantee rescue without imposing deleterious fitness costs, a feat that may be difficult to accomplish for essential genes requiring complex regulatory elements and networks not directly adjacent to the target gene. In contrast to the aforementioned drives, (i) HomeR relies on the endogenous promoter sequence of the target gene to facilitate rescue expression which significantly simplifies the design and ensures endogenous expression of the rescue using native regulatory machinery, (ii) creatively designed to target the 3’ end of the essential haplosufficient gene to limit the degree of recording required for the rescue, (iii) an exogenous 3’UTR to prevent recombination, and (iv) exploits BLM (Kandul et al., 2019b) by targeting an essential haplosufficient gene to convert recessive non-functional resistant alleles into dominant deleterious/lethal mutations that can get actively selected out of a population (**Fig. 1C, Fig. 6F, Fig. S1)**, four important features that should be incorporated into future population modification drives.

Results of three independent multi-generational population cage experiments initiating with 100% heterozygous populations (I.e. initial drive allele frequency of 50%; Cas9 allele frequency of 100%) indicate that NHEJ-induced loss-of-function (R2) alleles can persist for many generations. As expected, a single copy of the HomeR inserted at a haplosufficient gene provides sufficient rescue and complements the corresponding R2 allele. However, it was unexpected that up to 28% of sampled flies potentially harbor R2 *indels* that can persist for ten generations without being selected out (**Fig. 4A**). This high frequency of R2 alleles, for a HGD with nearly ∼100% transmission rate, suggest that these R2 alleles are likely induced from the paternal *wt* alleles by maternal carryover of Cas9/gRNA in zygotes harboring the maternal rescuing *Pol-γ35*^*HomeR1*^ allele (**Fig. 6F**). The maternal carryover can be a major source of both R2 and R1 resistance alleles, because it can possibly facilitate the cleavage and NHEJ repair of paternal *wt* allele before it comes into proximity with a maternal carrier allele to facilitate HDR (Adolfi et al., 2020; Champer et al., 2019; Gantz et al., 2015; Kandul et al., 2019a). Once R2 alleles are complemented by the *Pol-γ35*^*HomeR1*^ allele, it takes several generations for R2 alleles to combine as lethal homozygotes and be selected out from a population. The elimination of R2 alleles takes especially long time by HGDs targeting non-essential genes or genes whose knockouts do not cause complete lethality or sterility of homozygous carriers (Adolfi et al., 2020; Gantz et al., 2015) underscoring the importance of targeting essential haplosufficient genes. Therefore, to assess the stability of gene drives against accumulation of functional resistance alleles, its spread must be examined for several generations after a carrier frequency has reached 100%.

Our results are congruent with previous studies demonstrating reduced homing in *Drosophila* males (Chan et al., 2013, 2011; Windbichler et al., 2011). We tested multiple Cas9 lines supporting Cas9 expression in early and/or late germ cells with different levels of specificity, and have not achieved high levels of homing as reported in mosquito males, aka. >90% (Gantz et al., 2015; Kyrou et al., 2018). Achiasmatic meiosis in *Drosophila* males likely correlates with the weak activity of HDR pathway (Preston et al., 2006), which in turn results in inefficient homing in *Drosophila* males. Mosquito males have chiasmatic meiosis and recombination (Kitzmiller, 1976) that require active HDR machinery in primary spermatocytes, possibly contributing to efficient homing in mosquito males. Reduced homing efficacy in *Drosophila* males should be accounted for when designing HGDs in other species exhibiting achiasmatic meiosis, such as *D. suzukii*, an invasive fruit pest.

Our results indicate that functional resistance (R1) alleles can still be induced even when a conserved haplosufficient gene required for insect viability is targeted (**Fig. 4**). However, each identified in-frame, R1 allele changes at least one amino acid and thus may affect the fitness of its carrier preventing such alleles from accumulating at the expense of the drive. This incurring fitness cost likely slows down their accumulation and results in selection out of the population, in favor of the *Pol-γ35*^*HomeR1*^ alleles, over multiple generations (**Fig. 4**) again underscoring the importance of targeting an essential haplosufficient gene. Nevertheless, encoding additional gRNAs targeting the *wt* coding sequence of *Pol-γ35* downstream from the Cas9/gRNA cut site, which is re-coded in *Pol-γ35*^*HomeR1*^ alleles, into HomeR may further diminish the probability of inducing functional resistance alleles (Marshall et al., 2017).

Splitting HomeR into two genetic loci (*HomeR* and *Cas9*) integrated on different chromosomes serves as a molecular containment mechanism (**Fig. 5**). The *HomeR* element is able to home into *wt* alleles and bias its transmission. However, the *Cas9* element, which is inherited Mendelianly, is required for its homing. Therefore, the independent assortment of *Cas9* and *HomeR* limits the spread of *HomeR* and acts as a genetic ‘brake’ for HomeR propagation. The spread dynamic of split-HGDs resembles that of high-threshold drives and thus requires a high introduction rate for HomeR to spread into a local population and prevents its spread into neighboring populations (**Fig. 5**), which is an important feature for confining drive spread and may be necessary initial field testing of gene drives (Adelman et al., 2017; Akbari et al., 2015; Friedman et al., 2020; Kandul et al., 2019a; Li et al., 2020; Raban and Akbari, 2017; Raban et al., 2020). Moreover, if unintended consequences do arise, HomeR’s spread can be reversed by reintroduction of insects harboring *wt* alleles of the gene targeted by split-drive. Notwithstanding, if desired, HomeR could facilely be converted into a non-localized gene drive by incorporating the Cas9 into the Homer drive cassette. Taken together, the split-design of HomeR is safe localized gene drive technology that could be safely adopted and implemented for local control, and if a non-localized drive is desired for more wide scale spread, HomeR could be converted for that purpose too.

In sum, HomeR combines promising aspects of current population modification drives - confineablity, high transmission of HGD’s, and resilience to NHEJ generation of TA drives (**Fig. 6**). Modeling illustrates success of both design aspects in linked or split-drive form, demonstrating robust behavior over a range of parameter combinations (**Fig. 5**). This underscores its resilience to NHEJ alleles, overcoming a significant hurdle for current HGD designs. Given the simplicity of the HomeR design, it could be universally adapted to a wide range of species including human disease vectors in the future.

## Methods

### Selection of Cas9/gRNA target sites

We inserted a <u>Home</u>-and-<u>R</u>escue (HomeR) in *DNA Polymerase γ 35-kDa* (*Pol-γ35* or *PolγB*, CG33650). *Pol-γ35* is a haplosufficient gene required for insect viability: a lethal knockout can be rescued by a single functional copy. The highly conserved domain of *Pol-γ35* is located at the end of the coding sequence, which facilitates its re-coding (**Fig. 1**). We PCR amplified a 413-base fragment of the domain with 1073A.S1F and 1073A.S2R from multiple *Drosophila* strains (*w*^*1118*^, Canton S, Oregon R, *nos-Cas9 (Kandul et al*., *2019b)*) and used the consensus sequence along with the tool CHOPCHOP v2 (Labun et al., 2016) to choose two gRNA targets sites that minimize off-target cleavage.

### Design and assembly of genetic constructs

We used Gibson enzymatic assembly to build all genetic constructs (Gibson et al., 2009). To assemble both gRNA constructs, we used the previously described *sgRNA*^*Sxl*^ plasmid (Kandul et al., 2019b) (Addgene #112688) harboring the mini-*white* gene and attB docking site. We removed the fragment encompassing the U6.3 promoter and gRNA scaffold by AscI and SacII digestion, and we cloned it back as two fragments overlapping at a novel gRNA sequence (**Fig. 1A**). Both *U6*.*3-gRNA#1*^*Pol-γ35*^ and *U6*.*3-gRNA#2*^*Pol-γ35*^ plasmids targeting *Pol-γ35* are deposited at www.addgene.org (#159774 and #159675).

We assembled two *HomeR*^*Pol-γ35*^ constructs using two tested gRNAs (**Fig. 1 and 2**). Each *HomeR*^*Pol-γ3*5^ was built around a specific gRNA, with matching LHA and RHA: *HomeR1*^*Pol-γ35*^ harbored *U6*.*3-gRNA#1*^*Pol-γ35*^, and *HomeR2* had *U6*.*3-gRNA#2*^*Pol-γ35*^. We digested the *nos-Cas9* plasmid (Kandul et al., 2019b) (Addgene #112685) with AvrII and AscI, preserving the backbone containing the *piggyBac* left and right sequences that encompass the *Opie-dsRed-SV40* marker gene. The HomeR construct was assembled between *Opie-dsRed-SV40* and *piggyBacR* in three steps. First, we cloned the *U6*.*3-gRNA#1* or *#2* from the corresponding plasmid together with the *3xP3-eGFP-SV40* marker gene, to tag site-specific insertion of *GDe*. Then, we cloned three fragments: (1) LHA, which was amplified from the *Drosophila* genomic DNA; (2) the re-coded fragment downstream from the gRNA cut site, which was PCR amplified from the dePol-γ35 gBlock custom synthesized by IDT^®^ (**Table S1**); (3) the p10 3’UTR to provide robust expression (Pfeiffer et al., 2012) of the re-coded *Pol-γ35* rescue. Finally, we cloned RHA, which was PCR amplified from genomic DNA, corresponding to each specific gRNA cut site. Both *HomeR1*^*Pol-γ35*^ and *HomeR2*^*Pol-γ35*^ plasmids, targeting the *Pol-γ35* locus, are deposited at www.addgene.org (#159676 and #159677).

To assemble the three constructs for testis-specific Cas9 expression, we used a plasmid harboring the *hSpCas9-T2A-GFP*, the *Opie2-dsRed* transformation marker, and both *piggyBac* and attB-docking sites, which were previously used to establish Cas9 transgenic lines in *Aedes aegypti (Li et al*., *2017)* and *Drosophila melanogaster* (Kandul et al., 2019a, 2019b). We removed the *Ubiquitin 63E* promoter from the *ubiq-Cas9* plasmid (Addgene #112686) (Kandul et al., 2019b) by digesting it with SwaI at +27°C and then with NotI at +37°C, and cloned a promoter fragment amplified from the *Drosophila* genomic DNA. The *Drosophila exuperantia* (CG8994) 783-bp fragment (*exuL*) upstream of the *exuperantia* gene was amplified with ExuL.1F and ExuL.2R primers (**Table S1**) and cloned to assemble the *exuL-Cas9* plasmid. The *Rcd-1 related* (*Rcd1r*, CG9573) (Chan et al., 2013) and *β-Tubulin 85D (βTub) (Chan et al*., *2011; Michiels et al*., *1989)* promoters support early and late, respectively, testis-specific expression in *Drosophila* males. The 937-base-long fragment upstream of *Rcd1r* was amplified with 1095.C1F and 1095.C2R primers and cloned to assemble the *Rcd1r-Cas9* plasmid. The 481-base-long fragment upstream of *βTub* was amplified with βTub.1F and βTub.2R primers (**Table S1**) and cloned to build the *βTub-Cas9* plasmid. Three plasmids for testis-specific Cas9 expression are deposited at www.addgene.org (#159671 – 159773).

### Fly maintenance and transgenesis

Flies were maintained under standard conditions: 26°C with a 12H/12H light/dark cycle. Embryo injections were performed by Rainbow Transgenic Flies, Inc. We used φC31-mediated integration (Groth, 2004) to insert the *U6*.*3-gRNA#1* and *U6*.*3-g*RNA#2 constructs at the P{CaryP}attP1 site on the 2^nd^ chromosome (BDSC #8621), and the *exuL-Cas9, βTub-Cas9*, and *Rcd1r-Cas9* constructs at the PBac{y+-attP-3B}KV00033 on the 3^rd^ chromosome (BDSC #9750). Two methods were used to generate the site-specific insertion of *HomeR1*^*Pol-γ35*^ or *HomeR2*^*Pol-γ35*^ constructs at the *gRNA#1*^*Pol-γ35*^ or *gRNA#2*^*Pol-γ35*^ cut sites, respectively, inside the *Pol-γ35* gene via HDR. First, we injected the mixture of HomeR and helper *phsp-pBac*, carrying the piggyBac transposase (Handler and Harrell, 1999), plasmids (500 ng/µl and 250 ng/µl, respectively, in 30 µl) into *w*^*1118*^ embryos. Random insertions of *HomeR1*^*Pol-γ35*^ and *HomeR2*^*Pol-γ35*^, assessed by double (eye-specific GFP and body-specific dsRed) fluorescence (**Fig. 2D**), established with this injection were genetically crossed to *nos-Cas9/nos-Cas9* (BDSC #79004) (Kandul et al., 2019b) flies to “relocate” *HomeR1*^*Pol-γ35*^ or *HomeR2*^*Pol-γ35*^ to the corresponding gRNA cut site via Homology Assisted CRISPR Knock-in (HACK) (Lin and Potter, 2016). A few site-specific *Pol-γ35*^*HomeR1*^ and *Pol-γ35*^*HomeR2*^ lines tagged with only eye-specific GFP fluorescence were recovered. Second, we injected *HomeR1*^*Pol-γ35*^ or *HomeR2*^*Pol-γ35*^ plasmids directly into *nos-Cas9/nos-Cas9* (BDSC #79004) (Kandul et al., 2019b) embryos, generating multiple independent, site-specific insertions for each *Pol-γ35*^*HomeR*^ (**Fig. 2D**). Recovered transgenic lines were balanced on the 2^nd^ and 3^rd^ chromosomes using single-chromosome balancer lines (*w*^*1118*^; *CyO/sna*^*Sco*^ for II and *w*^*1118*^; *TM3, Sb*^*1*^*/TM6B, Tb*^*1*^ for III) or a double-chromosome balancer line (*w*^*1118*^; *CyO/Sp*; *Dr/TM6C, Sb*^*1*^).

We established three homozygous lines of *Pol-γ35*^*HomeR1*^ and *Pol-γ35*^*HomeR2*^ from independent insertion lines, and confirmed the precision of site-specific insertions by sequencing the borders between HomeR constructs and the *Drosophila* genome (**Fig. 2C**). The 1118-base-long fragment overlapping the left border was PCR amplified with 1076B.S9F and 1076B.S2R and was sequenced with 1076B.S3F and 1076B.S4R primers. The same-length fragment at the right border was amplified with 1073A.S1F and 1076B.S10R and was sequenced with 1076B.S7F and 1076B.S8R primers (**Supplementary Table 1**).

### Fly genetics and imaging

Flies were examined, scored, and imaged on a Leica M165FC fluorescent stereo microscope equipped with a Leica DMC2900 camera. We assessed the transmission rate of HomeR by following its eye-specific GFP fluorescence, while the inheritance of *Cas9* was tracked via body-specific dsRed fluorescence (**Fig. 2D, Fig. 4A**). All genetic crosses were done in fly vials using groups of ten males and ten females.

### *RNA*^*Pol-γ35*^ cleavage assay

To assess the cleavage efficiency of each gRNA targeting the C-terminal domain of *Pol-γ35*, we genetically crossed ten *w*^*1118*^; *U6*.*3-gRNA#1*^*Pol-γ35*^ or *w*^*1118*^; *U6*.*3-gRNA#2*^*Pol-γ35*^ homozygous males to ten *y*^*1*^, *Act5C-Cas9, w*^*1118*^, *Lig4 (X. Zhang et al*., *2014)* (BDSC #58492) homozygous females, and we scored the lethality of F_1_ males (**Fig. 1B**). The F_1_ males would then inherit the X chromosome from their mothers, expressing *U6*.*3-gRNA#1*^*Pol-γ35*^ or *U6*.*3-gRNA#2*^*Pol-γ35*^ with *Act5C-Cas9* in a *Lig4-*null genetic background, and this results in male lethality when a tested gRNA directs cleavage of the *Pol-γ35* locus. To assess the induced lethality in the *Lig4+/+* genetic background, we crossed ten *y*^*1*^, *Act5C-Cas9, w*^*1118*^ (BDSC #54590) (Port et al., 2014) flies to ten *U6*.*3-gRNA#1*^*Pol-γ35*^ flies in both directions, and scored survival of trans-heterozygous and heterozygous F_1_ progeny. To measure the Cas9/gRNA-directed cleavage of *Pol-γ35* by maternally deposited Cas9 protein in the *Lig4+* background, the same homozygous males were genetically crossed to *w*^*1118*^*/w*^*1118*^; *nos-Cas9/CyO* females (**Fig. 1C**), and the F_1_ progeny, harboring *U6*.*3-gRNA#1*^*Pol-γ35*^ or *U6*.*3-gRNA#2*^*Pol-γ35*^, were scored and compared to each other.

### Assessment of *Pol-γ35*^*HomeR*^ transmission rates

To compare transmission rates of *Pol-γ35*^*HomeR1*^ and *Pol-γ35*^*HomeR2*^, we first established trans-heterozygous parent flies by genetically crossing *Pol-γ35*^*HomeR1*^*/Pol-γ35*^*HomeR1*^; *+/+* or *Pol-γ35*^*HomeR2*^*/Pol-γ35*^*HomeR2*^; *+/+* females to *+/+*; *nos-Cas9/nos-Cas9* males. We then assessed the transmission rates by trans-heterozygous parent females and males crossed to *wt* flies. For controls, we estimated the transmission rates of *HomeR1*^*Pol-γ35*^ and *HomeR1*^*Pol-γ35*^ in the absence of Cas9, by heterozygous *Pol-γ35*^*HomeR1*^*/+* or *Pol-γ35*^*HomeR2*^*/+* females and males crossed to *wt* flies (**Fig. 3A**). To explore the effect of maternally deposited Cas9 protein on transmission of *Pol-γ35*^*HomeR1(Kandul et al*., *2019a)*^, we generated heterozygous *Pol-γ35*^*HomeR1*^*/CyO* embryos containing Cas9 protein deposited by *nos-Cas9/CyO* mothers and estimated the transmission of *Pol-γ35*^*HomeR1*^ by females and males raised from these embryos and crossed to *wt* flies. We tested five different Cas9 lines—supporting germline (*vas-Cas9)*, ubiquitous (*ubiq-Cas9, Act5C-Cas9*), and early (*exuL-cas9, Rcd1r-Cas9*) or late testes-specific expression (*βTub-Cas9*)—together with the strongest HomeR, *Pol-γ35*^*HomeR1*^. Ten trans-heterozygous females or males, generated by crossing homozygous *Pol-γ35*^*HomeR1*^ females to homozygous *Cas9* males, were genetically crossed to *wt* flies and the transmission of *Pol-γ35*^*HomeR1*^ was quantified in their F_1_ progeny (**Fig. 3C**).

### Egg hatching assay to assess the homing rate of *Pol-γ35*^*HomeR1*^

To identify the mechanism of the super-Mendelian transmission of *Pol-γ35*^*HomeR1*^, we assessed the percentage of F_1_ hatched eggs laid by trans-heterozygous *Pol-γ35*^*HomeR1*^*/+*; *nos-Cas9/+* females genetically crossed to *wt* males and compared it to those hatched from two types of heterozygous females: *Pol-γ35*^*HomeR1*^*/+*; *+/+* ♀ and *+/+*; *nos-Cas9/+* ♀ (**Fig. 3B**). We collected virgin females and aged them for three days inside food vials supplemented with a yeast paste, then five groups of 25 virgin females of each type were transferred into vials with fresh food containing 25 *wt* males and allowed to mate overnight (12 H) in the dark. Then, all males were removed from the vials, while females were transferred into small embryo collection cages (Genesee Scientific 59–100) with grape juice agar plates. After 12 H of egg laying, a batch of at least 200 laid eggs was counted for each sample group and incubated for 24 H at 26°C before the number of unhatched eggs was counted. Some fraction of hatched larvae for each test group was transferred into food vials to confirm that they would finish development.

### Accumulation of functional in-frame resistance alleles, *Pol-γ35*^*R1*^

To explore the generation and accumulation of functional resistance alleles induced by NHEJ, we initiated three drive populations by crossing 50 *+/+*; *nos-Cas9/nos-Cas9* females and 50 *Pol-γ35*^*HomeR1*^*/Pol-γ35*^*HomeR1*^; *nos-Cas9/nos-Cas9* (**Fig. 4A**) males in 0.3 L plastic bottles (VWR^®^ Drosophila Bottle 75813-110). Parent (P) flies were removed after six days, and their progeny were allowed to develop, eclose, and mate for 13–15 days. This established a 100% heterozygous *Pol-γ35*^*HomeR1*^*/+*; *nos-Cas9/+* population in the next generation (G_0_) (due to ∼100% transmission efficiency), with 50% allelic and 100% genotypic frequency of *Pol-γ35*^*HomeR1*^ in each bottle population. Each generation, around 250–350 emerged flies were anesthetized using CO_2_, and their genotypes with respect to *Pol-γ35*^*HomeR1*^ (presence or absence) were determined using the dominant eye-specific GFP marker. Then they were transferred to a fresh bottle and allowed to lay eggs for six days before removing them, and the cycle was repeated. Three populations were maintained in this way for eleven generations, which corresponds to ten generations of gene drive. Note that any fly scored without the *Pol-γ35*^*HomeR1*^ allele was transferred into a fresh bottle to ensure any *Pol-γ35* resistance or *wt* alleles could be passed to the next generation. We retrieved and froze the flies for genotyping only after six days to ensure sufficient time for breeding. We expected that the gRNA#1/Cas9-induced *Pol-γ35*^*R1*^ alleles that did not incur fitness costs would accumulate over a few generations and block the spread of the gene drive. However, as we did not find any fly without the *Pol-γ35*^*HomeR1*^ allele after G_3_, we stopped the population drives after ten generations of *Pol-γ35*^*HomeR1*^ homing in the heterozygous flies. We froze 60 flies after G_10_ for further sequence analysis.

### HomeR population replacement experiments

To assess the performance of HomeR GD experimentally, we established five drive and three control populations by seeding 50 homozygous *Pol-γ35*^*HomeR1*^ males and 50 *wt* males together with 100 *wt* virgin females in each a 0.3 L plastic bottle in the presence (homozygous *nos-Cas9* genetic background) or absence of *Cas9* (**Fig. 4C**). This ratio of *Pol-γ35*^*HomeR1*^ vs *Pol-γ35*^*WT*^ alleles resulted in the *Pol-γ35*^*HomeR1*^ introduction frequency of 25% in the parent generation. The discrete-generation populations were maintained and scored as described above. Each generation, around 250–350 emerged flies were anesthetized using CO_2_, and their genotypes with respect to *Pol-γ35*^*HomeR1*^ (presence or absence) were determined using the dominant eye-specific GFP marker. Then they were transferred to a fresh bottle and allowed to lay eggs for six days before removing them, and the cycle was repeated.

### Sequencing of induced resistance alleles

To analyze the molecular changes that caused functional in-frame (*R1*) and loss-of-function (LOF, *R2*) mutations in *Pol-γ35*, we PCR amplified the 232-base-long genomic region containing both *gRNA#1*^*Pol-γ35*^ and gRNA#2^Pol-γ35^ cut sites using 1073A.S3F and 1073A.S4R primers (**Table S1**). For PCR genotyping from a single fly, we followed the single-fly genomic DNA prep protocol (Kandul et al., 2019b). PCR amplicons were purified using the QIAquick PCR purification kit (QIAGEN), subcloned into the pCR™2.1-TOPO® plasmid (Thermofisher), and 5–7 clones were sequenced in both directions by Sanger sequencing at Retrogen^®^ and/or Genewiz^®^ to identify both alleles in a each fly. Sequence AB1 files were aligned against the corresponding *wt* sequence of *Pol-γ35* in SnapGene^®^ 4.

To explore the diversity of resistance alleles persisting after 10 generations of *Pol-γ35*^*HomeR1*^ in a 100% heterozygous population, we froze 60 flies (30 ♀ and 30 ♂), each harboring at least one copy of the dominant marker of *Pol-γ35*^*HomeR1*^, from each drive population after G_10_. Using these flies, we quantified any resistance and *wt* alleles remaining in the population via Illumina sequencing of heterogeneous PCR amplicons from the *Pol-γ35* locus. Note that PCR amplicons did not include the *Pol-γ35*^*HomeR1*^ allele due to its length (**Fig. 2A**). DNA was extracted using the DNeasy Blood and Tissue Kit (QIAGEN). To analyze heterogeneous PCR products, we used the Amplicon-EZ service by Genewiz^®^ and followed the Genewiz^®^ guidelines for sample preparation. In brief, Illumina adapters were added to the 1073A.S3F and 1073A.S4R primers to simplify the library preparation, PCR products were purified using QIAquick PCR purification kit (QIAGEN), around 50,000 one-direction reads covering the entire amplicon length were generated, and relative abundances of recovered SBS and *indel* alleles at the *gRNA#2*^*Pol-γ35*^ cut site were inferred using Galaxy tools (Afgan et al., 2018). Amplicon-EZ data from Genewiz^®^ were first uploaded to Galaxy.org. A quality control was performed using FASTQC. Sequence data were then paired and aligned against the *Pol-γ35*^*WT*^ sequence using Map with BWA-MEM under “Simple Illumina mode”. The SBS and *indel* alleles were detected using FreeBayes, with the parameter selection level set to “simple diploid calling”.

### Model fitting to cage experiment data

Empirical data from the *HomeR* population replacement experiments were used to parameterize a model of CRISPR-based homing gene drive including resistant allele formation. Model fitting was carried out for all five gene drive cage experiments using Markov chain Monte Carlo (MCMC) methods in which estimated parameters related to cleavage efficiencies in females and males, accurate HDR frequencies given cleavage in females and males, the proportion of resistant alleles that are in-frame and cost-free, and the fitness cost associated with having the *HomeR* system. We considered discrete generations, random mixing, and Mendelian inheritance rules at the gene drive locus, with the exception that for adults heterozygous for the homing allele (denoted by ‘H’) and wild-type allele (denoted by ‘W’), a proportion, *c*, of the W alleles are cleaved, while a proportion, 1-*c*, remain as W alleles. Of those that are cleaved, a proportion, *p*_*HDR*_, are subject to accurate HDR and become H alleles, while a proportion, (1-*p*_*HDR*_), become resistant alleles. Of those that become resistant alleles, a proportion, *p*_*RES*_, become in-frame, functional, cost-free resistant alleles (denoted by ‘R’), while the remainder, (1-*p*_*RES*_), become out-of-frame, non-functional, or otherwise costly resistant alleles (denoted by ‘B’). The values of *c* and *p*_*HDR*_ were allowed to vary depending on whether the HW individual is female or male. The fitness cost associated with the *HomeR* system, *s*_*H,F*_, was assumed to be female-specific. These considerations allowed us to calculate expected genotype frequencies in the next generation, and to explore the parameter values that maximize the likelihood of the experimental data. The model fitting framework is described in full in S1 Text of (Pham et al., 2019).

### Comparative modeling of gene drive systems

Comparative gene drive simulations were performed using a discrete-generation version of the Mosquito Gene Drive Explorer (MGDrivE) modeling framework (C. et al., 2020). The first generation was seeded with 200 adults, 75% wild-type and 25% heterozygous for each gene drive, split equally between sexes. At each generation, adult females mate with males, thereby obtaining a composite mated genotype (their own, and that of their mate) with mate choice following a multinomial distribution determined by adult male genotype frequencies. Egg production by mated adult females then follows a Poisson distribution, proportional to the genotype-specific lifetime fecundity of the adult female. Offspring genotype follows a multinomial distribution informed by the composite mated female genotype and the inheritance pattern of the gene drive system. Sex distribution of offspring follows a binomial distribution, assuming equal probability for each sex. Female and male adults from each generation are then sampled equally to seed the next generation, with a sample size of 200 individuals (100 female and 100 male), following a multivariate hypergeometric distribution. 25 repetitions were run for each drive in the trace plots (**Fig. 5A and 5C**) and 100 repetitions were run for each parameter combination in the heatmaps (**Fig. 5B and 5D**).

The inheritance pattern is captured by the “inheritance cube” module of MGDrivE. ClvR and TARE constructs were implemented to match their published descriptions (Champer et al., 2020a; Oberhofer et al., 2020a, 2019). HomeR and HGD were implemented as one or two-locus systems following equivalent inheritance rules. When Cas9 and gRNAs co-occur in the same individual, wild-type alleles are cleaved at a rate *c*_*F*_ (*c*_*M*_) (female-(male-) specific cleavage), with 1-*c*_*F*_ (1-*c*_*M*_) remaining wild-type. Given cleavage, successful HDR occurs at a rate *ch*_*F*_ (*ch*_*M*_), with 1-*ch*_*F*_ (1-*ch*_*M*_) alleles undergoing some form of NHEJ. Of these, a proportion, *cr*_*F*_ (*cr*_*M*_), are in-frame NHEJ alleles, while the remainder, 1-*cr*_*F*_ (1-*cr*_*M*_), are LOF alleles. Maternal carryover (maternal deposition, or maternal perdurance) was modeled to occur in zygotes of mothers having both Cas9 and gRNAs, impacting a proportion, *d*_*F*_, of zygotes. Of the wild-type alleles in impacted zygotes, a proportion, *dr*_*F*_, become in-frame NHEJ alleles, while the remainder, 1-*dr*_*F*_, become LOF alleles. These inheritance rules apply to both HomeR and HGD, with differing fitness costs.

ClvR (Oberhofer et al., 2020a, 2019) was modeled using a 99% cleavage rate in female and male germ cells, as well as in embryos from maternal carryover. For two-locus ClvR, the two loci were assumed to undergo independent assortment (>=50cM separation), as was assumed for all two-locus systems in this analysis. For both configurations, it was assumed that 0.1% of cleaved alleles were converted to functional resistance alleles (R1 type), and the rest became LOF alleles (R2 type). In addition to the 50% egg-hatching reduction due to the non-homing drive (**Fig. 6A-B**), an additional 5% reduction in fecundity was applied to females that harbored Cas9. For consistency, TARE, HGD, and HomeR (for ideal parameters) also used a cleavage rate of 99% in females and males, though TARE demonstrated lower maternal carryover (Champer et al., 2020a), and was modeled with 95% cleavage. HGD and HomeR (for ideal parameters), which rely on HDR, were simulated with 90% HDR rates in females and males. Cleaved alleles that did not undergo HDR were assumed to be R1 alleles with proportion 0.5%, and R2 LOF alleles the remainder of the time. TARE and HomeR were also modeled with a small (5%) fitness reduction, applied as a reduction of female fecundity. Since a HGD does not provide a rescue for a knocked out targeted gene, its carriers demonstrate higher fitness costs, and were assigned a 20% fitness reduction with the assumption that the HGD is inserting into a non-lethal gene that imposes a low/moderate fitness cost. Experimentally-derived parameters for HomeR differed from ideal parameters in two ways: i) there was no HDR in males (although cleavage remained the same), and ii) 1% of NHEJ-repaired *wt* alleles were converted into R1 alleles (c.f. 0.5% for the ideal case). All simulations were performed, analyzed, and plotted in R (“Website,” n.d.) (R Core Team 2017). Code is available upon request.

### Statistical analysis

Statistical analysis was performed in JMP 8.0.2 by SAS Institute Inc., and graphs were constructed in Prism 8.4.1 for MacOS by GraphPad Software LLC. At least three biological replicates were used to generate statistical means for comparison. *P* values were calculated using a two-sample Student’s *t*-test with equal or unequal variance.

### Gene Drive safety measures

All crosses using gene drive genetics were performed in accordance to a protocol approved by the Institutional Biosafety Committee at UCSD, in which full gene drive experiments were performed in a high-security ACL2 barrier facility and split-drive experiments were performed in an ACL1 insectary in plastic vials that were autoclaved prior to being discarded, in accordance with currently suggested guidelines for the laboratory confinement of gene drive systems (Akbari et al., 2015; National Academies of Sciences, Engineering, and Medicine et al., 2016).

### Ethical conduct of research

We have complied with all relevant ethical regulations for animal testing and research and conformed to the UCSD institutionally approved biological use authorization protocol (BUA #R2401).

### Data and Reagent Availability

All data are represented fully within the tables and figures. The *U6*.*3-gRNA#1*^*Pol-γ35*^, *U6*.*3-gRNA#2*^*Pol-γ35*^, *HomeR1*^*Pol-γ35*^, *HomeR2*^*Pol-γ35*^, *exuL-Cas9, Rcd1r-Cas9*, and *βTub-Cas9* plasmids and corresponding fly lines are deposited at Addgene.org (159671–159677) and the Bloomington Drosophila Stock Center, respectively.

### Disclosures

O.S.A is a founder of Agragene, Inc., has an equity interest, and serves on the company’s Scientific Advisory Board. N.P.K is a consultant for Agragene. All other authors declare no competing interests.

## Author Contributions

N.P.K and O.S.A. conceived the idea and designed experiments. N.P.K and J.L. engineered plasmids, and performed all molecular and genetic experiments. J.B.B and J.M.M performed mathematical modeling. All authors analyzed the data, contributed to the writing of the manuscript, and approved the final manuscript.

### Acknowledgments

This work was supported in part by funding from a DARPA Safe Genes Program Grant (HR0011-17-2-0047), and NIH awards (R21RAI149161A, R01AI151004, DP2AI152071) awarded to O.S.A. The generation of the *ExuL* promoter was done by O.S.A. while at Caltech working with Bruce A. Hay.

## Supplementary Information

**Supplementary Figure 1.**
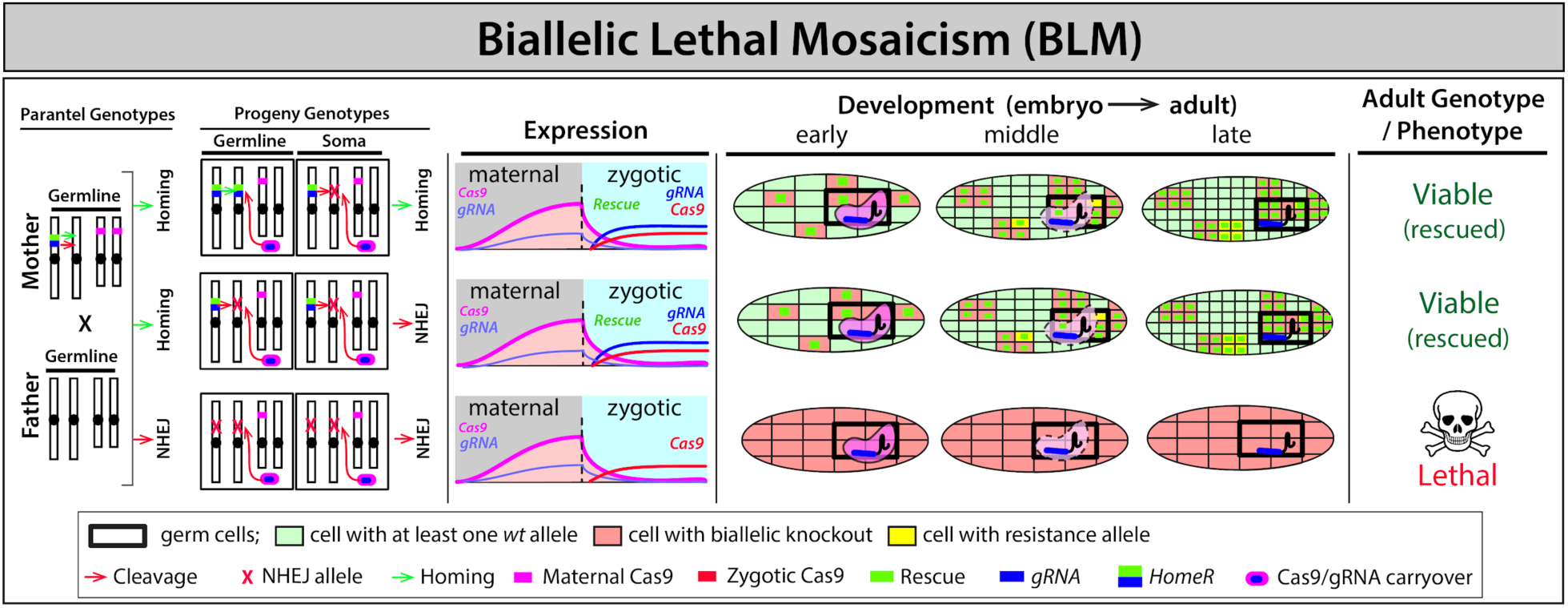
Mechanism of biallelic lethal mosaicism (BLM). Maternal carryover of Cas9/gRNA complexes contributes to RNA-guided dominant biallelic knockouts (both paternal and maternal alleles) of an essential target gene throughout development thereby converting recessive non-functional resistant alleles into dominant deleterious/lethal mutations that can get negatively selected out of a population (Kandul et al., 2019b). Individuals that inherit the HomeR drive are protected through expression of a dominant recoded protected copy of the haplosufficient essential target gene (rescue) and are therefore viable/fertile, while individuals that inherit two disrupted alleles are dead (i.e. target gene is recessive lethal).

**Supplementary Figure 2.**
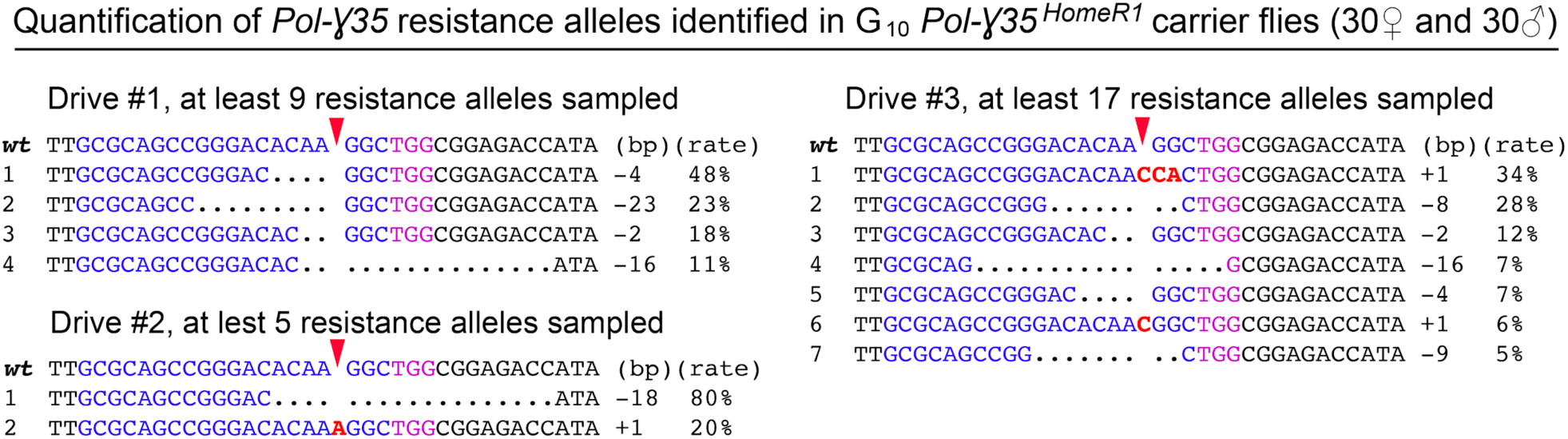
Quantification of *Pol-γ35* resistance alleles sampled after ten generations of *Pol-γ35*^*HomeR1*^ homing. Resistance alleles, persisting for ten generations of cleavage and homing, were sampled from sixty flies chosen randomly harboring at least one *Pol-γ35*^*HomeR1*^ allele and were quantified using Illumina® sequencing. For each gene drive, nearly 50,000 amplicons of *Pol-γ35* alleles (150K total), which did not carry the 2.5 kb insert of the *HomeR1*, were sequenced and used to estimate the minimum number of sampled resistance alleles. Note that both functional in-frame resistance alleles (*R1*.*1* and *R1*.*2*) identified at earlier generations (**Fig. 4B**) were not sampled after generation 10. Two novel in-frame resistance alleles (−18 bp and −9 bp) resulted in deletions of 6 and/or 3 amino acids and would not be good rescues. The sequence of *gRNA#1*^*Pol-γ35*^ is highlighted in blue, and its PAM sequence is in purple. Red arrows depict Cas9/gRNA cut sites. Base insertions and amino acid changes are in red.

**Supplementary Table 1**. Target sequence of gRNA and primers used in this study.

**Supplementary Data 1**. Cleavage assay of two *gRNAs*^*Pol-γ35*^ with *Act5C-Cas9* in the *Lig4Δ* genetic background.

**Supplementary Data 2**. Cleavage assay of two *gRNAs*^*Pol-γ35*^ with *nos-Cas9*.

**Supplementary Data 3**. Transmission rate of *Pol-γ35*^*HomeR1*^ or *Pol-γ35*^*HomeR1*^ in conjunction with different Cas9 lines.

**Supplementary Data 4**. Assessment of embryonic lethality induced by *Pol-γ35*^*HomeR1*^*/+; nos-Cas9/+* females.

**Supplementary Data 5**. Spread of resistance alleles in *Pol-γ35*^*HomeR1*^/+; *nos-Cas9/nos-Cas9* flies over ten generations.

